# Abundant clock proteins point to missing molecular regulation in the plant circadian clock

**DOI:** 10.1101/2024.09.03.609973

**Authors:** Uriel Urquiza-García, Nacho Molina, Karen J. Halliday, Andrew J. Millar

**Author notes:** email: Uriel Urquiza-García, Nacho Molina, Karen Halliday, Andrew Millar.

## Abstract

Understanding the biochemistry behind whole-organism traits such as flowering time is a longstanding challenge, where mathematical models are critical. Very few models of plant gene circuits use the absolute units required for comparison to biochemical data. We refactor two detailed models of the plant circadian clock from relative to absolute units. Using absolute RNA quantification, a simple model predicted abundant clock protein levels in *Arabidopsis thaliana*, up to 100,000 proteins per cell. NanoLUC reporter protein fusions validated the predicted levels of clock proteins *in vivo*. Recalibrating the detailed models to these protein levels estimated their DNA-binding dissociation constants (*K_d_*). We estimate the same *K_d_* from multiple results *in vitro*, extending the method to any promoter sequence. The detailed models simulated the *K_d_* range estimated from LUX DNA-binding *in vitro* but departed from the data for CCA1 binding, pointing to further circadian mechanisms. Our analytical and experimental methods should transfer to understand other plant gene regulatory networks, potentially including the natural sequence variation that contributes to evolutionary adaptation.

## Introduction

Circadian clocks are intracellular regulators that control temporal gene expression patterns and hence metabolism, physiology and behaviour, from sleep/wake cycles in mammals to flowering in plants (Bass & Takahashi, 2010; Bendix *et al*, 2015; Millar, 2016). Clock genes are rarely essential but appropriate alleles can confer a competitive advantage (Ouyang *et al*, 1998; Dodd *et al*, 2005), have been repeatedly selected during crop domestication (Bendix *et al*, 2015; Muller *et al*, 2015) and are implicated in human disease, including cancers, metabolic and mental health (Roenneberg *et al*, 2022). Systems biology uses models to link these organismal traits to molecular pathways, in order to understand and potentially to engineer circadian functions (Clark *et al*, 2020; Chew *et al*, 2022). Plant biologists have a particular opportunity to connect molecular understanding to the rich tradition of crop science models (Thomas, 2007; Marshall-Colon *et al*, 2017; Hammer *et al*, 2019), alongside large-scale plant phenomics data (Tardieu *et al*, 2017). This study focuses on the clock gene circuit in the laboratory model plant *Arabidopsis thaliana*, as the non-transcriptional timing mechanism in this species has not been characterised (Edgar *et al*, 2012).

The Arabidopsis clock circuit comprises a dozen genes with tightly-interlinked feedbacks that are sufficient to generate 24-hour rhythmicity in mathematical models (see below). To simplify, dawn-expressed transcription factors *LATE ELONGATED HYPOCOTYL* (*LHY*) and *CIRCADIAN CLOCK-ASSOCIATED 1* (*CCA1*) inhibit the expression of evening genes such as *GIGANTEA* (*GI*), EARLY FLOWERING genes (*ELF3* and *ELF4*) and LUX ARTHYHMO (*LUX*) by directly binding to their promoter regions through the ‘Evening Element’ target sequence (Adams *et al*, 2018; Harmer & Kay, 2005; Kamioka *et al*, 2016; Nagel *et al*, 2015). *LHY* and *CCA1* expression is ended by the binding of repressors from the *PSEUDO-RESPONSE REGULATOR* (*PRR*) gene family, which are expressed in the day, in sequence *PRR9*, *PRR7*, *PRR5* and *TOC1* (*TIMING OF CAB2 EXPRESSION 1*) (Nakamichi *et al*, 2010; Huang *et al*, 2012; Gendron *et al*, 2012). Falling LHY and CCA1 protein levels allow the expression of ELF3, ELF4 and LUX proteins that form an “Evening Complex” in the early subjective night (Nusinow *et al*, 2011). *Via* the LUX subunit (also known as PHYTOCLOCK1), the complex binds to and represses the expression of *TOC1* and several evening genes (Helfer *et al*, 2011; Silva *et al*, 2016), while the PRR proteins degrade, allowing *LHY* and *CCA1* expression in the late night to start the cycle anew. The pace of this repressor-based circuit is modified by transcriptional activators (Perez-Garcia *et al*, 2015; Shalit-Kaneh *et al*, 2018; Urquiza-García & Millar, 2021) and by post-translational regulation, including from light input pathways, which entrain the clock to ensure that rhythmic activities occur at appropriate phases relative to the external, day/night cycle (Millar, 2016).

A series of mathematical models has represented the biochemistry of the clock gene circuit with increasing detail in differential equations (Pokhilko *et al*, 2012, 2013; Fogelmark & Troein, 2014; Urquiza-García & Millar, 2021), while other models used simpler versions (Dalchau *et al*, 2011; De Caluwé *et al*, 2016; Foo *et al*, 2020; Greenwood *et al*, 2022). The models have previously predicted new molecular clock components and interactions (reviewed in Bujdoso & Davis, 2013), and explained some operating principles of the clock mechanism (Akman *et al*, 2008; Edwards *et al*, 2010; Gould *et al*, 2013; Rand, 2008). Modelling the circadian control of downstream pathways (Seaton *et al*, 2015) has allowed us to bridge the genotype-phenotype gap, linking molecular regulation to whole-organism traits (Chew *et al*, 2022). However, the detailed models have two general limitations. First, they used real time units but arbitrary mass units, so the values of many biochemical-kinetic parameters could not be validated against biochemical data, such as synthesis rates or binding constants. One exception recently introduced absolute RNA levels (Urquiza-García & Millar, 2021). Second, the genes in these models are functional units with no internal structure, so they cannot directly represent genetic variation at the level of genome sequence. Here, we refactor the Arabidopsis clock model to represent plant clock protein levels in absolute units of protein copies per cell, introduce tractable methods that facilitate validation against biochemical and *in vivo* data, and extend one method to include genome sequence directly.

Clock protein numbers have been both measured and modelled in the fungus *Neurospora crassa* (Merrow *et al*, 1997; Smolen *et al*, 2003), in mammalian cells (Gabriel *et al*, 2021; Koch *et al*, 2022; Kramer *et al*, 2020; Narumi *et al*, 2016; Smyllie *et al*, 2016) and in the cyanobacterium *Synechococcus elongatus* (Chew *et al*, 2018; Kitayama *et al*, 2003). The absolute numbers of proteins directly constrain their possible biochemical activities (Kim & Forger, 2012) and dynamic behaviour (Leise *et al*, 2012; Chew *et al*, 2018; Gould *et al*, 2018), refining our understanding of clock mechanisms. Consider, for example, a clock transcription factor that rhythmically binds to the promoters of target genes. A protein with a high DNA-binding affinity (low dissociation constant, *K_d_*) cannot be such a rhythmic regulator, if it is active at concentrations well above the *K_d_* at all circadian phases, binding to its target sites throughout the day and night. Hence the levels of the clock proteins constrain their possible dissociation constants and *vice versa*, if the protein is to function as expected, in this case as expected in a mathematical model. The plant clock models include this mutual constraint but their *K_d_* values cannot be compared to measured binding data, because the models’ *K_d_* values have arbitrary units.

### Clock protein levels as a Fermi problem

We approach this question as a “Fermi problem” (Phillips & Milo, 2009), combining the available data from diverse sources to give coarse predictions for the numbers of plant clock protein molecules per cell in *Arabidopsis thaliana* (Figure 1). We rescale the detailed clock models for these protein levels, which also returns simulated *K_d_* values for their DNA binding in absolute units. The model predictions are compared with data-driven estimates of these dissociation constants, using an approach that allows promoter sequence data to inform the clock models. Lastly, we use reporter fusion proteins to measure the clock protein levels, largely validating our initial predictions, in plant extracts and by *in vivo* imaging.

**Figure 1.**
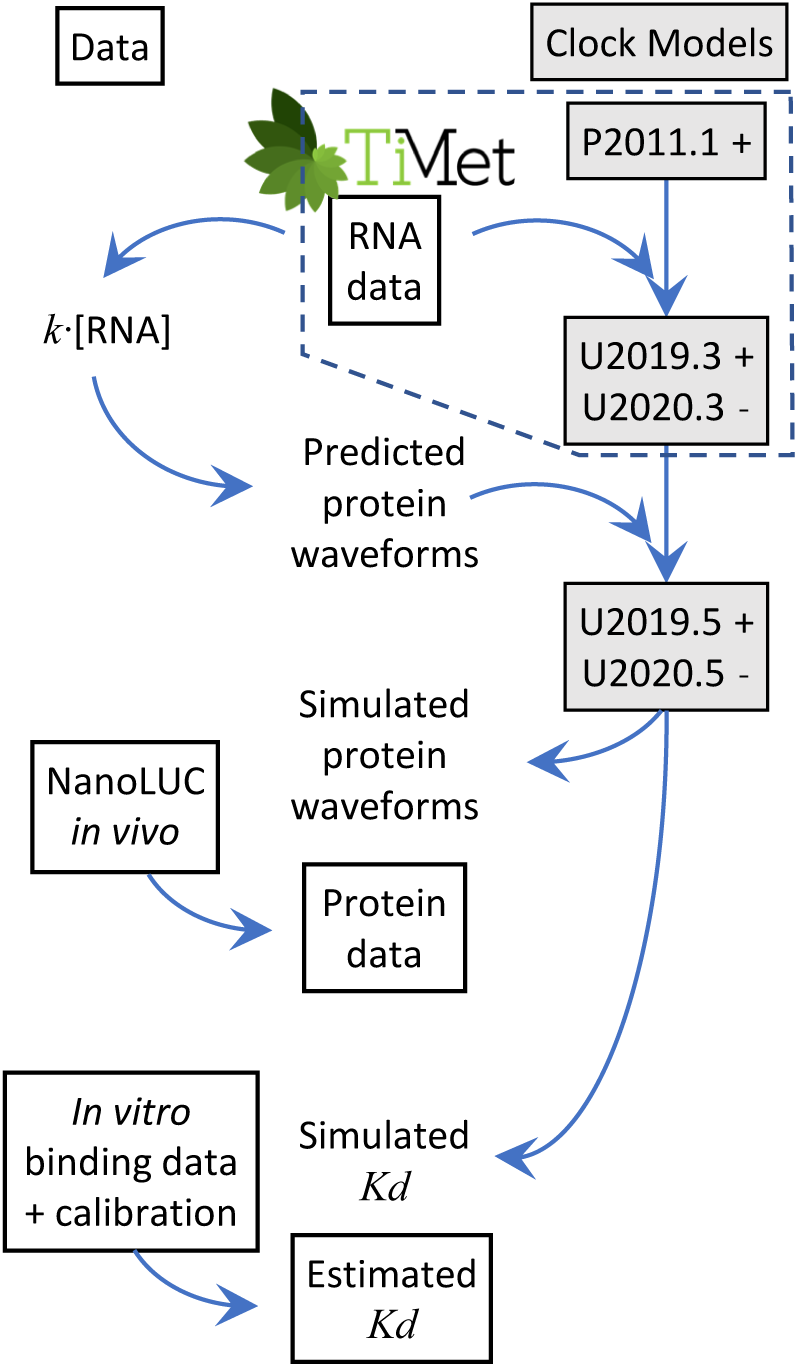
Model calibration and testing. Models of the Arabidopsis circadian clock (shaded boxes), with PRR activation (+) or inhibition (-), were calibrated to match experimental data (open boxes). Absolute RNA levels from the TiMet project were previously used to calibrate the U2019 and U2020 models (dashed outline; Urquiza-García & Millar, 2021). Here, a simple protein model (k.[RNA]) uses the same RNA data to estimate absolute protein levels and recalibrate new versions of each model. The models first simulate detailed protein waveforms that are tested against absolute protein data from NanoLUC reporter genes in transgenic plants. Second, the models predict transcription factor dissociation constants for DNA-binding (Kd) in vivo. These are tested against the constants estimated in vitro from surface plasmon resonance data, protein-binding microarrays, genome sequences transformed by a binding energy matrix, ChIP-seq and nuclear volume data.

## Results

### Predicting clock proteins levels from mRNA data in absolute units

We extended an earlier method that was applied to predict metabolic enzyme levels in moles per gramme fresh weight (Piques *et al*, 2009), in order to predict the levels of clock proteins in units of molecules per cell. Timeseries of mRNA transcript levels for multiple clock proteins were previously measured in molecules per cell (Figure 2a), using calibrated qRT-PCR assays (Flis *et al*, 2015). If the relevant translation and degradation rates are known, protein levels can be estimated from these RNA data using a simple, data-driven model,

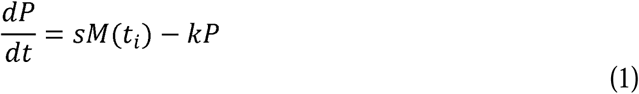

**Figure 2.**
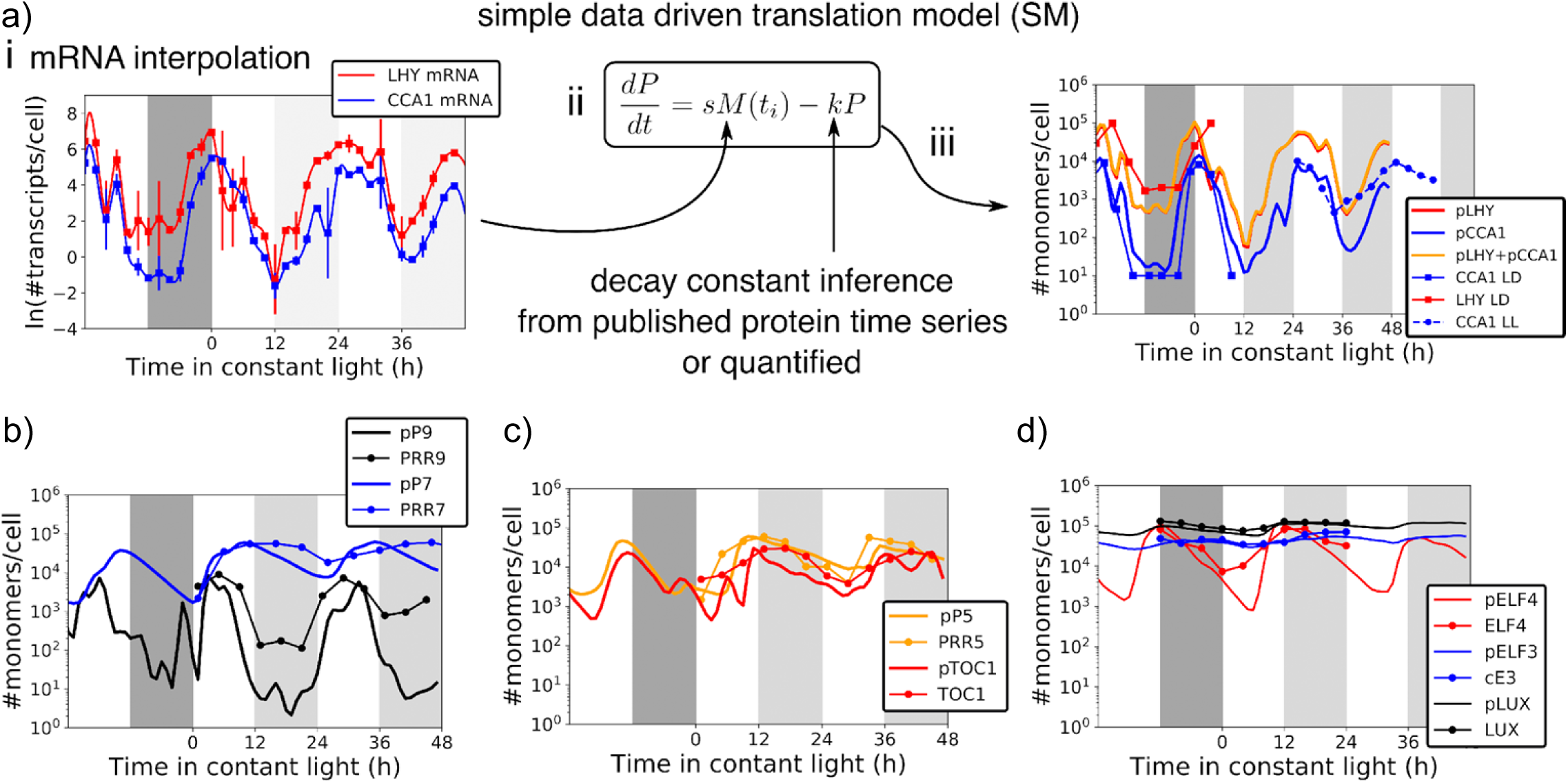
Predicting clock protein levelsfrom RNA data. a) i. mRNA timeseries data from the TiMet project (Flis *et al*, 2015) was log transformed and interpolated with a cubic spline (*LHY* mRNA, red; *CCA1* mRNA, blue). ii, interpolated RNA data *M(t)* was the input to the simple model. The translation rate *s* was calculated and protein decay constants *k* were derived from literature, as described in Results. c-f) Protein timeseries predicted from the simple model (solid lines without markers, protein names with prefix p) in absolute units are plotted with protein timeseries from the literature (with markers), for (a.iii) LHY and CCA1, (b) PRR9 and PRR7, (c) PRR5 and TOC1 and (d) ELF3, ELF4 and LUX. Literature data in arbitrary units were rescaled to the maximum value of the model prediction within each timeseries, to compare protein waveforms. a.iii-d) have the same logarithmic scale.

Where *P* stands for protein amount, *s* for translation rate, *M(t_i_)* mRNA transcript level in absolute units at time *t=i* and *k* is the decay constant of the protein of interest (Figure 2b). We follow past work (Piques *et al*, 2009) in estimating the translation rate *s* for each protein, from a ribosome elongation rate measured in mammalian cells, corrected for the lower growth temperature of Arabidopsis plants, to give 3 codons/s, using the measured ribosome density of 6.6 ribosomes per kb mRNA transcript from *E. coli*, and lengths of open reading frames from the TAIR10 genome annotation (see Supplementary Information). Effective degradation rates *k* were taken from measured protein decay curves for LHY and CCA1 proteins (Hansen *et al*, 2017; Song & Carré, 2005), and for PRR7, PRR9, PRR5 and TOC1 proteins (Farre & Kay, 2007; Ito *et al*, 2007; Kiba *et al*, 2007; Mas *et al*, 2003). Decay timeseries data for the Evening Complex proteins has not been reported, so the effective rates were estimated by fitting (see Supplementary Information) to profiles derived from published protein gel blot images (Nusinow *et al*, 2011). We also digitised full timeseries profiles under LD and/or LL for the above-mentioned proteins from the literature, using the same approaches. Light strongly destabilises some clock proteins, so separate, light and dark estimates were used for some degradation rates in the data-driven model (Supplementary Table 1), whereas the smaller effect of light on translation rate was ignored (Kim *et al*, 2003; Piques *et al*, 2009).

The predicted protein timeseries closely followed the mRNA profiles (compare Figures 2a, 2c). Table 1 shows that the predicted levels for the clock proteins of interest peaked at around 100,000 protein copies per cell and fell to trough levels in the range of 1000-5000 protein copies per cell. PRR9 had a lower predicted level due to its low RNA level (Figure 2d). The predicted amplitude of protein regulation changed in constant light conditions (LL) compared to light:dark cycles, for example from a 32-fold amplitude of predicted PRR7 protein rhythmicity in LD to 5-fold amplitude under LL (Table 1). This reflected the lower amplitude of mRNA rhythms observed in general under LL (Flis *et al*, 2015) (Figure 2d). The timing of the simple model’s predicted waveforms closely matched the peaks of the protein gel blot profiles (Figure 2c-f), though PRR7 had a broader peak during the subjective night in the data. Amplitude information is hard to derive by digitising gel blot images from the literature.

**Table 1.** Protein levels predicted by the simple model, simulated by the full models and observed in extracts or *in vivo*. Peak and trough levels are given as the protein copy number per cell, along with relative amplitude (fold change). Levels under 12L:12D cycles (LD) are all from ZT0-24h of the relevant timeseries. Levels in constant light conditions (LL) are from the cycle indicated (ZT LL). LL/LD ratio is the relative amplitude in LL compared to LD. Simulated levels (Sim.) combine all pools of each protein. Levels were observed (Obs.) using NanoLUC fusion reporters (NL) either in plant extracts (*in vitro* assay) or *in vivo*. Values for LUX are included for completeness; * measurements for LUX-NL are preliminary due to partial complementation by the fusion protein.

| Conditions: |  | LD |  |  | ZT LL | LL |  |  | LL/LD ratio | plot |
| --- | --- | --- | --- | --- | --- | --- | --- | --- | --- | --- |
| Protein | Model or data | peak | trough | fold-change |  | peak | trough | fold-change |  |  |
| CCA1+LHY | Pred. | 89506 | 460 | 194.6 | 12-34 | 58786 | 449 | 130.9 | 0.67 | Figure 2a)iii |
| CCA1/LHY | Sim. U2019.5 | 103803 | 10041 | 10.3 | 12-34 | 53061 | 7825 | 6.8 | 0.66 | Figure 3b LHY+CCA1 panel |
| CCA1/LHY | Sim. U2020.5 | 114645 | 8410 | 13.6 | 12-34 | 37379 | 6326 | 5.9 | 0.43 | Supplementary Figure 1a |
| LHY-NL | Obs. *in vitro* | 121118 | 1431 | 84.6 | 12-34 | 188712 | 3484 | 54.2 | 0.64 | Figure 6a |
| CCA1-NL | Obs. *in vitro* | 182067 | 4682 | 38.9 | 12-34 | 93877 | 2816 | 33.3 | 0.86 | Figure 6a |
| PRR7 | Pred. | 37341 | 1613 | 23.2 | 24-48 | 59200 | 7605 | 7.8 | 0.34 | Figure 2b |
| PRR7 | Sim. U2019.5 | 42004 | 3124 | 13.4 | 24-48 | 28623 | 5558 | 5.1 | 0.38 | Figure 3b PRR7 panel |
| PRR7 | Sim. U2020.5 | 54589 | 152 | 359.1 | 24-48 | 13885 | 2829 | 4.9 | 0.01 | Supplementary Figure 1b |
| PRR7-NL | Obs. *in vitro* | 81208 | 4761 | 17.1 | 24-48 | 55881 | 3875 | 14.4 | 0.85 | Figure 6b |
| TOC1 | Pred. | 23313 | 493 | 47.3 | 12-34 | 29981 | 1877 | 16.0 | 0.34 | Figure 2c |
| TOC1 | Sim. U2019.5 | 33978 | 3872 | 8.8 | 12-34 | 48752 | 3221 | 15.1 | 1.72 | Figure 3b TOC1 panel |
| TOC1 | Sim. U2020.5 | 14759 | 2639 | 5.6 | 12-34 | 14443 | 2342 | 6.2 | 1.10 | Supplementary Figure 1c |
| TOC1-NL | Obs. *in vitro* | 44482 | 3523 | 12.6 | 12-34 | 67995 | 3523 | 19.3 | 1.53 | Figure 6c |
| ELF3 | Pred. | 44868 | 26280 | 1.7 | 24-48 | 57426 | 29879 | 1.9 | 1.13 | Figure 2d |
| ELF3 | Sim. U2019.5 | 19813 | 2783 | 7.1 | 24-48 | 28035 | 7212 | 3.9 | 0.55 | Figure 3b ELF3 panel |
| ELF3 | Sim. U2020.5 | 11319 | 516 | 21.9 | 24-48 | 14705 | 2827 | 5.2 | 0.24 | Supplementary Figure 1d |
| ELF3-NL | Obs. *in vivo* | 41798 | 24897 | 1.7 | 24-48 | 28877 | 17892 | 1.6 | 0.96 | Figure 6f |
| LUX | Pred. | 99830 | 59001 | 1.7 | 24-48 | 122351 | 57535 | 2.13 | 1.26 | Figure 2d |
| LUX | Sim. U2019.5 | 167943 | 15502 | 10.8 | 24-48 | 165069 | 12957 | 12.74 | 1.18 | Figure 3b LUX panel |
| LUX | Sim. U2020.5 | 145379 | 6949 | 20.9 | 24-48 | 138629 | 10261 | 13.51 | 0.65 | Supplementary Figure 1g |
| LUX-NL* | Obs. *in vivo* * | 4455 | 1577 | 2.8 | 24-48 | 6355 | 1807 | 3.52 | 1.24 | Supplementary figure 4f |
| LUX-NL* | Obs. *in vitro* * | 596 | 379 | 1.6 | 24-48 | 882 | 362 | 2.4 | 1.55 | Supplementary figure 5c |

### Detailed models with absolute protein copy numbers

Our detailed models U2019.3 and U2020.3 rescaled the clock’s RNA variables into absolute units, in two versions of the clock gene circuit (Urquiza-García & Millar, 2021). That work used the same RNA timeseries data from the TiMet project as the simple model, above. We could now rescale these mechanistic models from protein levels in arbitrary units to the absolute units predicted in Table 1 (equations in Supplementary Files 1, 2; for SBML model files, please see Data Availability). The scale of all protein variables in the models is uniquely defined by a subset of proteins, the combined LHY/CCA1 protein, PRR7 and LUX. The PRR proteins and TOC1 function additively as transcriptional repressors in the model, so rescaling any one of them sets the scale of the other three. The stoichiometry of the Evening Complex also couples the scale of its components but of these, only LUX binds to DNA and provides DNA-binding affinity data to compare with the model prediction. The three protein scaling factors were estimated numerically by fitting the model simulations to the protein timeseries predicted from the simple model (Figure 3a; LHY/CCA1 was fitted to total predicted LHY+CCA1; see Methods).

**Figure 3.**
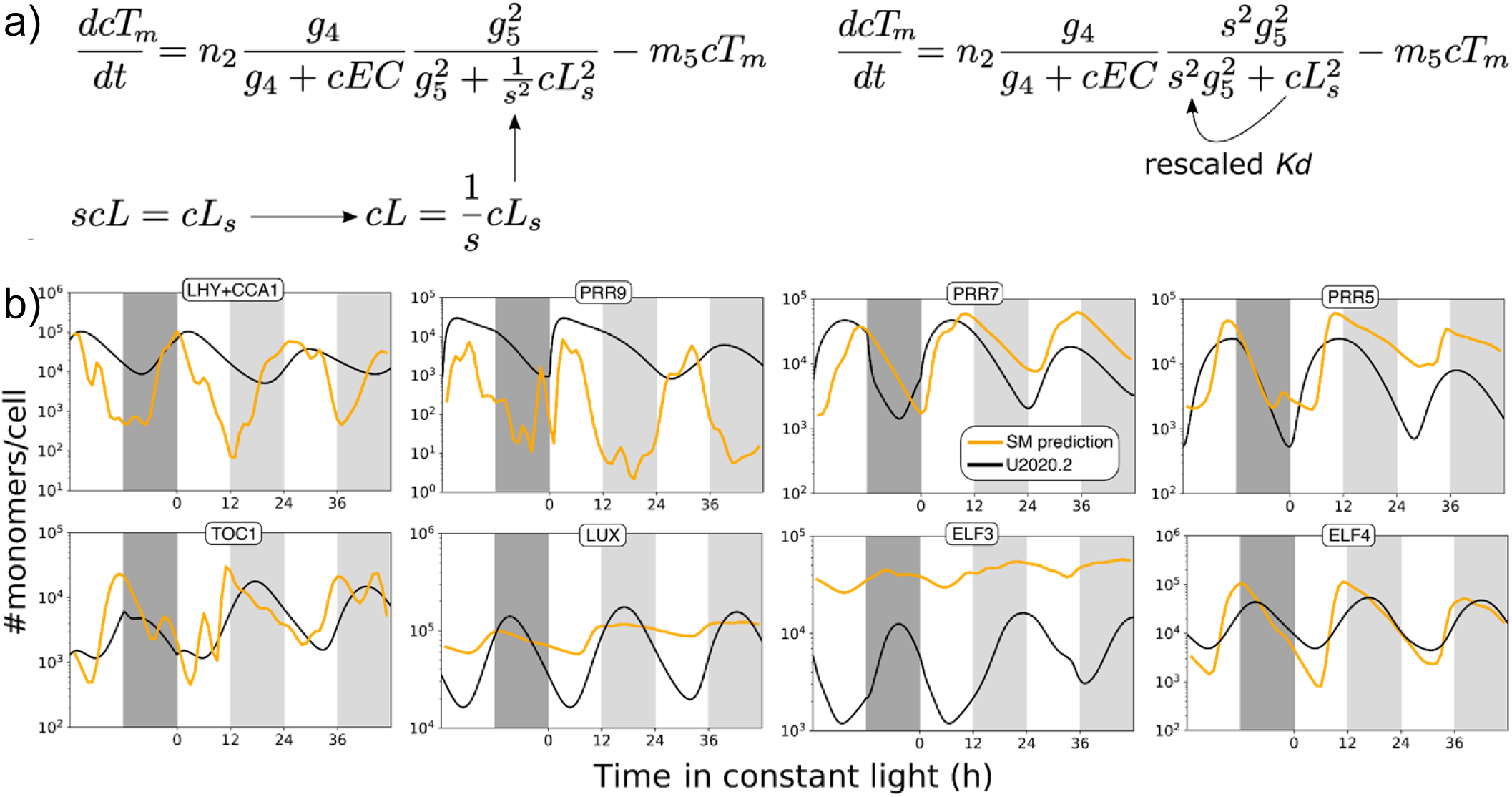
Rescaling protein levels in the mechanistic models. Scaling factors were introduced into the protein equations to form models U2019.4 and U2020.4. In (a), variable *cL* (LHY/CCA protein) in arbitrary units is replaced by *cL_s_* in molecules per cell, by introducing scaling parameter *s*. This example shows the equation for *TOC1* mRNA *cT_m_*, where *cEC* is the Evening Complex repressor copy number; *g, m* and *n* parameters are listed in Supplementary Tables 2 and 3. In U2019.5 and U2020.5, all parameters are re-scaled, so *g_5_* takes the value *s*·*g_5_*. (b) Fitting the simulated proteins to the protein waveforms predicted from the simple model (orange lines) estimated the best-fit values of the scaling factors. The resulting models U2019.4 (b) and U2020.4 (Supplementary Figure 1) simulate protein dynamics in molecules per cell (black lines). Simulated ELF3 protein remained below the predicted levels. The models were entrained to ten 12L:12D cycles prior to the interval plotted. Plots are scaled differently for each protein; Figure 2 shows all predicted proteins on the same scale.

The resulting models U2019.4 and U2020.4 retain the same dynamic behaviour as the parent models but simulate the clock proteins in units of molecules per cell. The version number U2019.4 indicates a model with the U2019 regulatory circuit but different parameter values from U2019.3, for example. Comparing the simple, data-driven model and the rescaled, full model, we found similar protein levels and dynamics in most cases, with expected departures where the regulation differed between the simple and full models (Figure 3b; Supplementary Figure 1). The full models’ predicted amplitudes for the PRR9 or LHY/CCA1 protein rhythms did not fully match the very high amplitudes of RNA regulation that drive the simple model, for example. More strikingly, the best-fit rescaling in both U2019.4 and U2020.4 left the mean level of ELF3 protein an order of magnitude lower than predicted from the RNA data (Figure 3b; see Discussion). The simple model’s predicted protein levels alone were potentially informative but the detailed models link the protein levels to more detailed timeseries simulations, and also to the proteins’ functions in DNA binding.

### Comparing model predictions to measured DNA-binding affinities

Models U2019.5 and U2020.5 propagated the re-scaling from the scaling factors alone to all the model parameters (Supplementary Tables 2 and 3), including the clock proteins’ dissociation constants for DNA-binding (*K_d_*). The rescaled models predicted relatively high *K_d_* values (low binding affinities) for LHY/CCA1 in the 100 nM range (Table 2, Supplementary Table 4). These are within the 0.167 - 700 nM range of measured dissociation constants that were previously collated from the literature for plant DNA-binding proteins (Millar *et al*, 2019). *K_d_* values that fall within the range of protein levels simulated in the models indicate a substantial change in binding over the circadian cycle, for example in the binding of LHY/CCA1 to the *PRR9* promoter (Figure 4). However, these simulated *K_d_* values are two orders of magnitude higher than the lowest *K_d_* measured by Surface Plasmon Resonance, 1.44 +/- 0.2 nM, for bacterially-expressed LHY/CCA1 heterodimers binding to a consensus Evening Element sequence (O’Neill *et al*, 2011). 1.44 nM corresponds to ∼100 proteins per nucleus. One further analysis recalibrated the simulated *K_d_* to account for ∼1000 genome-wide CCA1 binding sites that are absent from the model (please see Discussion).

**Figure 4.**
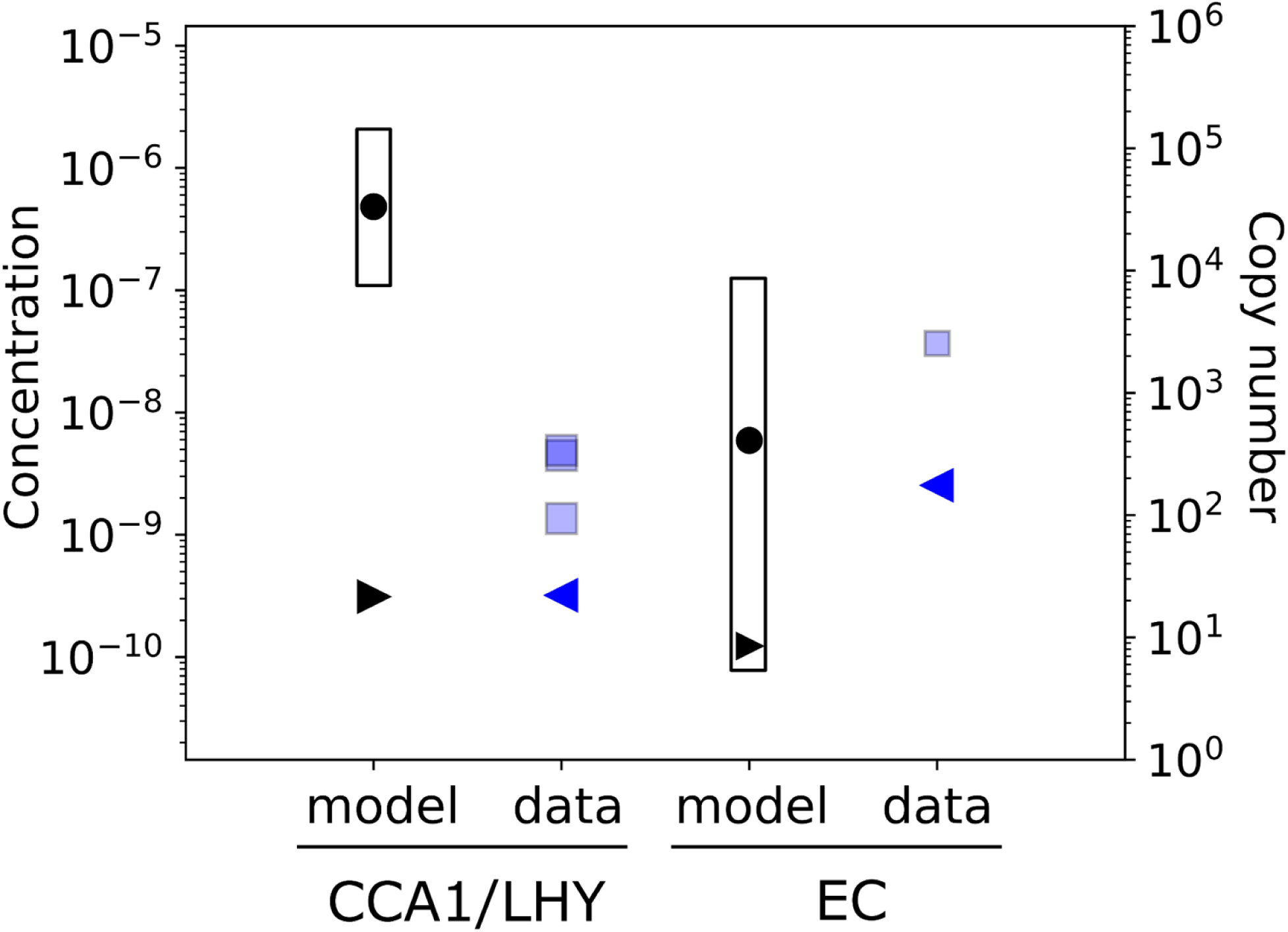
Protein function in DNA binding, in simulations and data. The plots show the dynamic range of the simulated CCA1/LHY protein (*cL*; left) or simulated Evening Complex (EC, *cEC*; right) in model U2019.5 under 12L:12D cycles (open bar), in units of molecules per cell (right axis) or nuclear concentration (left axis). The simulated *K_d_*’s for binding of each protein to *PRR9* (parameters *g9* and *g8*, respectively) fall within that range (black point). One possible calibration for the number of ChIP-seq targets in the genome yields the black triangles (pointing right; see Discussion). Blue squares represent the *K_d_* measured *in vitro* for CCA1, LHY, or mixed CCA1 and LHY (forming heterodimers), or for the LUX DNA-binding domain in the EC, each tested on its cognate consensus binding sequence from the *PRR9* promoter. Extending these *K_d_*’s to the *PRR9* upstream region gives the blue triangles (pointing left). The dynamic ranges and *K_d_*’s for other genes are shown in Tables 1, 2 and Supplementary Table 4.

**Table 2.** Clock protein dissociation constants for DNA-binding, from models or data. Examples of simulated *K_d_*’s in model U2019.5 are listed by the protein variable, such as cEC for the Evening Complex, the regulated variable, such as *cE4m* for *ELF4* mRNA, and the name of the *K_d_* parameter, such as *g2*. *K_d_* /ChIP peaks, a possible calibration of the simulated *K_d_* by the number of genomic regions bound in ChIP-seq data, see Discussion. Kd’s derived using EMA from experimental data are listed by protein and target gene. N/A, ChIP-seq data indicate no binding. The activator *cLmod* presumably corresponds to RVE8 and other proteins, for which no binding data are available.

| Model regulator | Model target | parameter | Sim. Kd (copy number/cell) | Sim. Kd (nM) | Kd / ChIP peaks (nM) | Clock protein | Target gene | EMA Kd (nM) |
| --- | --- | --- | --- | --- | --- | --- | --- | --- |
| *cEC* | *cE4m* | *g2* | 596 | 8.6 | 0.18 | LUX | *ELF4* | 1.68 |
| *cEC* | *cGm* | *g14* | 206 | 3.0 | 0.06 | LUX | *GI* | 2.49 |
| *cEC* | *cP9m* | *g8* | 408 | 5.9 | 0.12 | LUX | *PRR9* | 2.54 |
| *cEC* | *cTm* | *g4* | 451 | 6.5 | 0.14 | LUX | *TOC1* | 3.17 |
| *cL* | *cE3m* | *g16* | 28141 | 408 | 0.26 | CCA1 | *ELF3* | N/A |
| *cL* | *cE4m* | *g6* | 24508 | 355 | 0.23 | CCA1 | *ELF4* | 0.25 |
| *cL* | *cGm* | *g15* | 42247 | 612 | 0.40 | CCA1 | *GI* | 0.58 |
| *cL* | *cLUXm* | *g6* | 24508 | 355 | 0.23 | CCA1 | *LUX* | 0.34 |
| *cL* | *cP9m* | *g9* | 33342 | 483 | 0.31 | CCA1 | *PRR9* | 0.32 |
| *cL* | *cTm* | *g5* | 20979 | 304 | 0.20 | CCA1 | *TOC1* | 0.41 |
| *cL+cLmod* | *cP7m* | *g10* | 67505 | 978 | 0.63 | CCA1 | *PRR7* | 0.32 |
| *cLmod* | *cP5m* | *g12* | 16905 | 245 | 0.16 | RVE8 etc. | *PRR5* |  |

The detailed models predict low nM values for LUX binding to its promoter targets as part of the Evening Complex (EC; Table 2). These *K_d_*’s lie within the range of simulated EC levels (Figure 4), consistent with phase-specific binding of the EC in the model. The simulated *K_d_* are below or overlapping the 6.5 to 43 nM range of *K_d_*’s measured for the LUX DNA-binding domain on consensus target sequences (Silva *et al*, 2016, 2020). Hence the models’ simulated *K_d_*’s were much closer to the values measured *in vitro* for LUX, where only the LUX protein in the EC can bind its targets, than the equivalent data for LHY and CCA1. However, the simulated EC copy number ranges from tens to thousands per cell (Figure 4), at least an order of magnitude less than total LUX. No such *K_d_* data were available for PRR proteins.

Among the differences between the rescaled models and the measured *K_d_*’s (see also Discussion; Supplementary Table 5), the model *K_d_*’s necessarily reflect the binding of the clock protein over the entire target gene, because the model’s genes have no internal structure. In contrast, the *K_d_*’s measured *in vitro* are for individual, high-affinity, consensus sequences. A target gene might have several exact copies of the consensus sequence or none. We therefore developed an approach to extrapolate the dissociation constants for arbitrary promoter sequences, using large-scale data that is available for *Arabidopsis thaliana*.

### Extending the approach to promoter sequences

O’Neill *et al*. (2011) measured the *K_d_* of LHY and/or CCA1 binding *in vitro* to only three target sequences, two high-affinity sites and a “non-binding”, mutated sequence (Harmer & Kay, 2005). Similar data for LUX were obtained by Electromobility Shift Assay (EMSA) (Silva *et al*, 2016, 2020). Fortunately, Protein Binding Microarray (PBM) assays (Figure 5a; Berger & Bulyk, 2009) have been applied to test the binding of CCA1 and LUX to all possible 8-mer sequences (Franco-Zorrilla *et al*, 2014; Helfer *et al*, 2011). Unfortunately, their results in Enrichment score (E-score) units were not directly comparable to biophysical *K_d_* measurements. Comparing the CCA1 E-scores for the three target sequences tested by O’Neill *et al*., their *K_d_*‘s suggested a simple, linear relationship between the microarray results and the biophysical assays (Figure 5d). Applying this linear regression to the microarray dataset would assign tentative dissociation constants for CCA1 binding to any 8- mer sequence. We performed a similar analysis for LUX, where the dissociation constants for several consensus binding sequences were obtained by Electromobility Shift Assay (EMSA) (Silva *et al*, 2016, 2020). The linear relationship observed there also (Figure 5e) supported our approach of using PBM data as a proxy for *K_d_* (see Supplementary Information).

**Figure 5.**
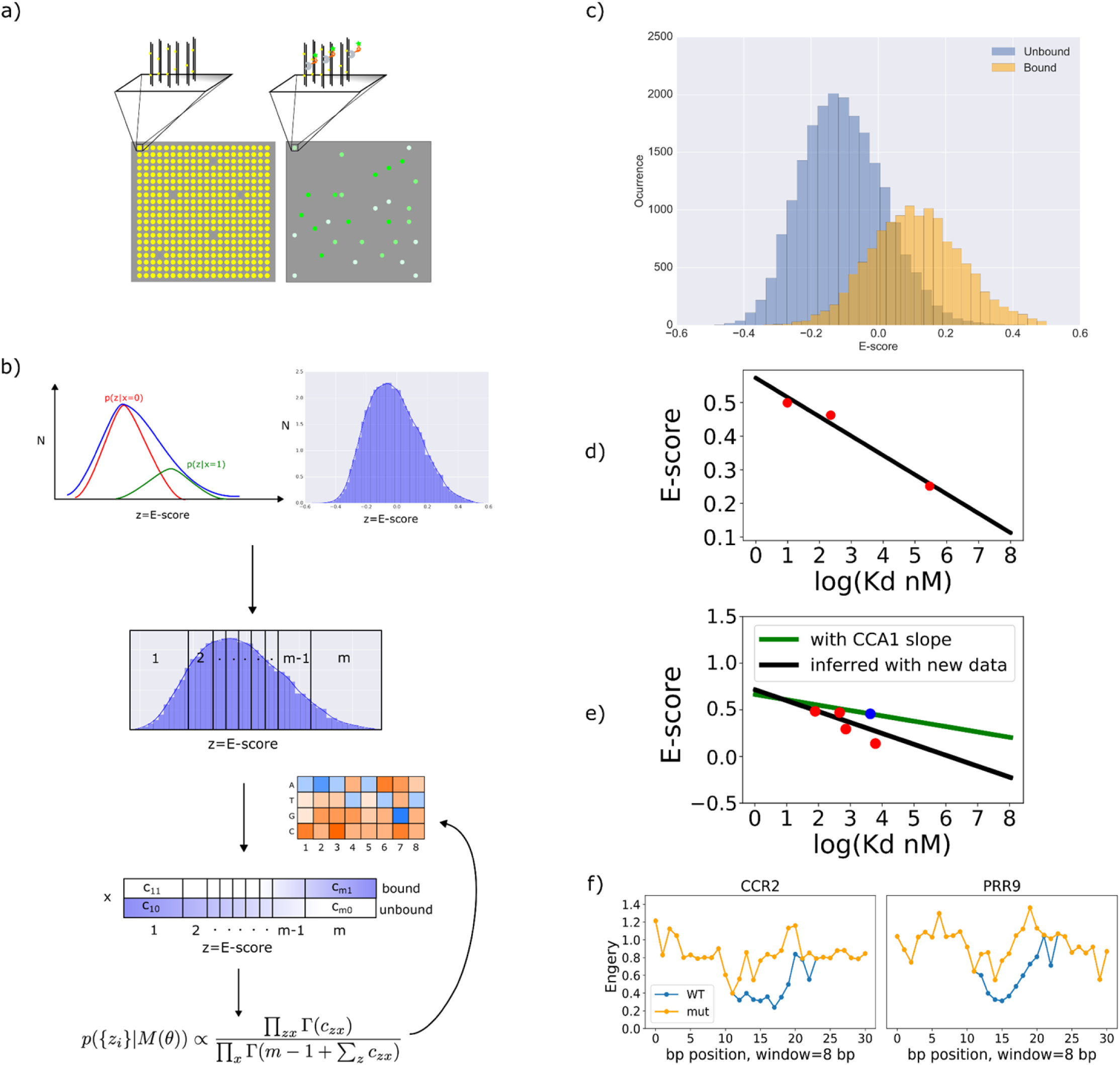
Applying measured *Kd*’s to any binding sequence. (a) Schematic of Protein Binding Microarray (PBM, left) and the fluorescence data from binding tagged proteins (green, right), which are quantified (b) as an E-score for each 8-mer DNA binding sequence (blue line). The overlapping distributions of E-scores from unbound (red line) and bound (green line) sequences can be estimated. The Error Model Averaging algorithm (EMA; Kinney *et al*, 2007) considers the E-score distribution in bins. EMA infers a best-fit matrix of binding energies, for each nucleotide at each position of the 8-mer (rectangle) and classifies bound (c, yellow) and unbound sequences. The PBM data included the sequences used to measure the binding *K_d_* (in nM units) *in vitro* for CCA1 (d) and LUX (e). Only these sequences have binding data (red points) both in E-score and nM units, defining linear relationships (solid lines) that can recalibrate the E-score of any other sequence to a *K_d_* in nM. (e) also plots a potential alternative calibration (green line), applying the slope of the CCA1 calibration line to a single *K_d_* for LUX (blue point; Silva *et al*, 2016). (f) shows the predicted energy of CCA1 binding to an 8-basepair sequence window, in the promoters of *CCR2* (left) or *PRR9* (right) around canonical binding sites (blue line) or mutated versions (orange line).

A technical concern arose, because the “non-binding”, mutated version of the high-affinity CCA1 target sequence had a measured E-score around 0.25, indicating greater CCA1 binding than to many other sequences. Experimental error in the microarray assay is expected to further confound signals, where weak binding and non-binding sequences overlap in this part of the E-score distribution (Figure 5b). To avoid introducing an arbitrary threshold between binding and non-binding sequences, the biophysical Error Model Averaged (EMA) approach (Kinney *et al*, 2007; Barnes *et al*, 2019) was applied (Figure 5c) to deconvolve the distributions of bound and unbound sequences in the E-scores (Figure 5d), using a binding energy matrix inferred for the 8 base-positions x 4 bases in each possible target sequence on the microarray (see Supplementary Information).

Finally, the sequences predicted to bind CCA1 (or LUX) were identified upstream of each gene in the model. The total *K_d_* was calculated by adding the contribution of their constituent 8-mers, such that this *K_d_* estimate is directly informed by the promoter sequences (Figure 5f). The approach might be refined to account for mutual interference among DNA-binding proteins, nucleosomes (Sullivan *et al*, 2014; Zhang *et al*, 2015) and other factors in future (Supplementary Table 5). The result is a *K_d_* for each target gene derived from direct DNA-binding data for CCA1 and LUX, which can be compared to the *K_d_* values simulated independently by the models in absolute units (Table 2, Supplementary Table 4).

These total *K_d_* values were typically an order of magnitude lower than the values reported for the consensus sequences alone (Figure 4, Table 2). For LUX, the resulting estimated *K_d_*’s of 1.3-3.2 nM (median 2.5nM) were all within the models’ 0.3-8.6 nM range of simulated *K_d_*’s (median 5.3nM), suggesting remarkable agreement between data and model. For CCA1, these lower *K_d_* values for the target clock genes were still further from the models’ simulated *K_d_* values (Table 2, Supplementary Table 4). The *K_d_* values in U2019.5 and U2020.5 depend on the protein levels predicted by the simple model. If the predicted LHY and CCA1 levels were too high, the full models would simulate incorrectly-high *K_d_* values, so we tested those predictions by measuring clock protein levels *in vivo*.

### Absolute quantification of clock proteins using NanoLUC reporter fusions

Antibodies are available for few of the plant clock proteins, and their low abundance has made them challenging to quantify by mass spectrometry, though each of these approaches holds promise for the future. Our recent implementation of the small, ostracod-derived, Nano-luciferase (NanoLUC) reporter protein in transgenic plants (Urquiza-García & Millar, 2019) suggested an alternative strategy, using calibrated luciferase assays to test the levels of reporter fusions to the clock proteins. To retain alternative splicing and other post-transcriptional regulation (James *et al*, 2018), we inserted a dual-epitope-tagged, NanoLUC-3xFLAG-10xHis reporter (NL3F10H) into genomic clones of the clock genes *LHY, CCA1, PRR7, TOC1, LUX* and *ELF3*, encoding translational fusions between each clock protein and a C-terminal NanoLUC (Supplementary Figure 2; Supplementary Tables 6, 7). After testing reporter activity in transient, protoplast transfections (Supplementary Figure 3), stable transgenic lines were generated in the background of the cognate mutant (or double mutant) for each clock gene. Lines were selected by quantitative complementation of the clock-mutant phenotype (Supplementary Figure 4), implying that the fusion protein’s expression level was functionally similar to the wild-type protein. Complementation was tested using the circadian period of a transcriptional, firefly luciferase (LUC+) reporter transgene and/or a physiological, hypocotyl elongation assay (see Supplementary Information; Supplementary Table 8).

The selected, transgenic lines were grown under light:dark cycles (LD) for 21 days and sampled every 2h, followed by a transfer to constant light (LL), in conditions similar to the TiMet RNA timeseries study. Known amounts of purified, recombinant NanoLUC were used to calibrate *in vitro* assays of whole-cell extracts from a measured fresh weight of transgenic plant material, yielding rhythmic timeseries data for LHY, CCA1, PRR7 and TOC1, in units of molecules per cell (Figure 6a-6c). The measured levels of the fusion proteins were very similar to those predicted by the simple model, with peak protein expression of LHY, CCA1 and PRR7 around 100,000 molecules per cell under LD conditions and trough levels in the low to mid 1000’s (Table 1). TOC1 peaked only slightly lower, at 44,000 molecules per cell. These direct experimental measurements validated the simple model’s predicted protein levels.

**Figure 6.**
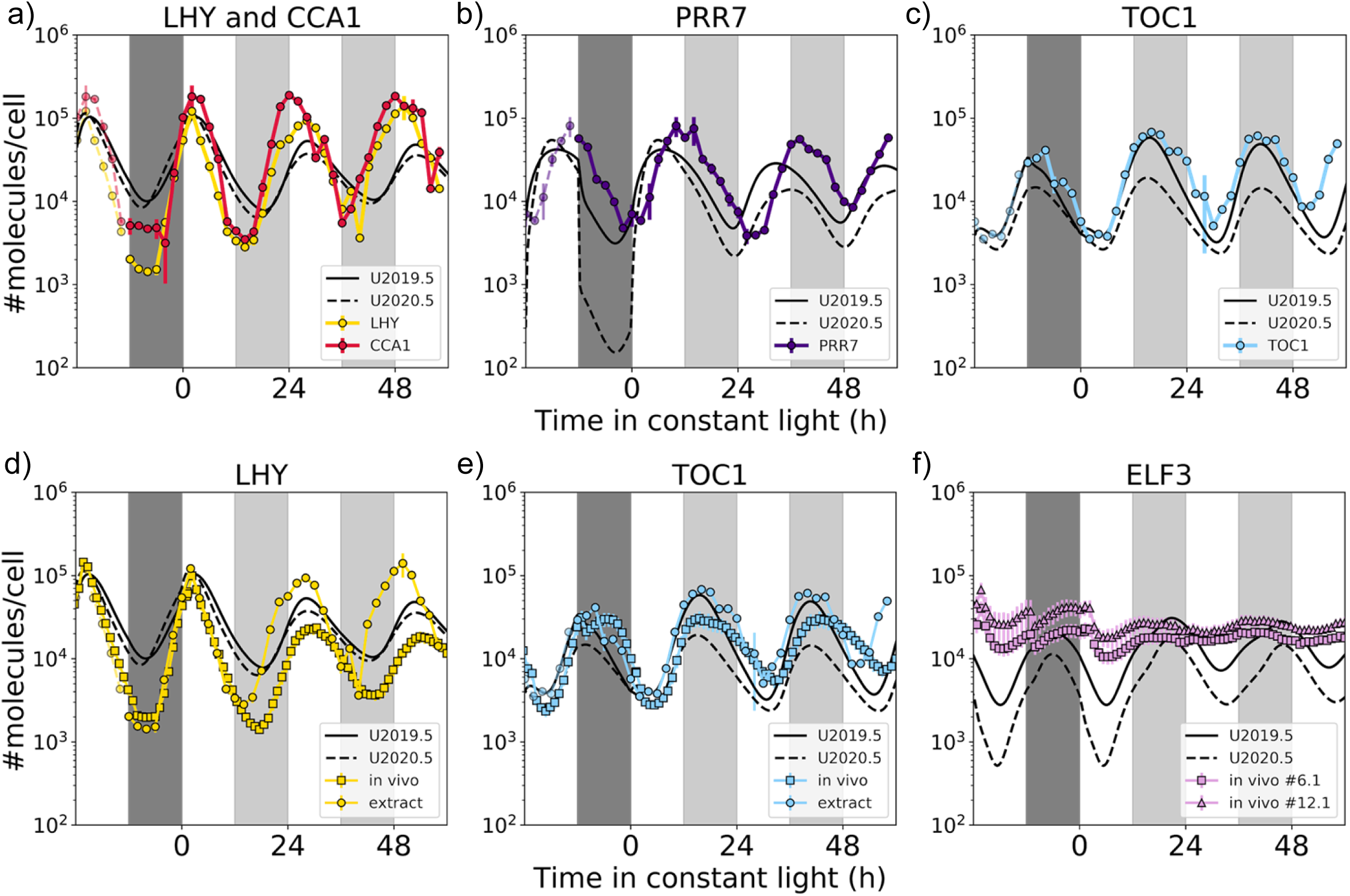
Absolute protein quantification verifies the mass scale of the clock models. (a-c) Reporter protein levels were measured in calibrated NanoLUC assays, for fusions to (a) LHY (yellow) and CCA1 (red), (b) PRR7 and (c) TOC1, in extracts of plants harvested every 2h from dusk under a 12L:12D cycle followed by 60h constant light from time 0h. Measured protein levels closely match the simulated levels from models U2019.5 (solid black line) and U2020.5 (dashed black line). Plants were grown for 21 days in 12L:12D before sampling started. Data from 0-10h are double-plotted at −24 to −14h (dashed lines). Protein data are means of duplicate biological samples, each of 5 plants, with technical triplicate assays; error bar = 1 SEM. (d-f) *in vivo* reporter assays suggest ELF3 levels. Seedlings were grown in micro-well plates for 10 days in 12:12D and furimazine substrate was added to each well. NanoLUC activity was measured hourly, in seedlings carrying LHY (d), TOC1 (e) and ELF3 (f) reporters, under one further 12L:12D cycle followed by constant light in an automated luminometer. The falling trend due to furimazine decay was removed from the *in vivo* signals (Supplementary Figure 6). A single scaling value matched *in vivo* data for LHY (d) and TOC1 (e) to the cognate signals from extracts (a, c). The same scaling factor was applied to the data from two ELF3 reporter lines (f), suggesting the levels of ELF3 protein *in vivo*. Light interval, white background; dark interval, dark grey shading; anticipated dark interval during constant light, light grey shading.

The LUX fusion line tested in extracts showed much lower expression levels (mid-100s of molecules per cell; Supplementary Figure 5c), and contrasted with the strong expression of a closely-related BOA protein reporter (Urquiza-García & Millar, 2019). Further testing showed that many LUX fusion lines restored the rhythmic expression of our screening marker in the *lux-2* mutant host, but failed to complement its long-hypocotyl phenotype (Supplementary Figure 4e). A second LUX protein reporter line that complemented both mutant phenotypes was therefore selected and tested by *in vivo* luminometry, along with a reporter for ELF3, as described below.

### High-resolution timeseries data compares measured and modelled plant clock proteins

The timeseries profiles of the fusion transgenes were very similar to earlier gel blot data and slightly lagged the RNA timeseries, confirming that the reporters were appropriately regulated, with peak levels of LHY at ZT2, CCA1 at ZT2-4, PRR7 at ZT10 and TOC1 at ZT16 under LD conditions (Figure 6a-6c). LHY and CCA1 protein levels remained low during an extended trough of 6-10h in the late day to early night, while PRR7 and TOC1 levels peaked. Low signal variation among the biological triplicate samples suggested that the technical data quality from the *in vitro* NanoLUC assays was comparable to the TiMet qRT-PCR timeseries, allowing detailed interpretation of the protein timeseries.

Comparing the protein data to the predicted protein levels in the full model (Figure 6), the LHY and CCA1 protein data are similar in timing to the simulated CCA1/LHY and overlap in levels, though the data has a longer, lower protein trough in LD. The measured PRR7 waveform is shifted about 2h later than the model under LD, but with similar protein levels to the models under LL. PRR7 showed less protein degradation in the dark than predicted in the U2020.5 model; the protein data were closer to the U2019.5 profile. Phase plane diagrams compared the TiMet timeseries for *LHY* and *CCA1* mRNA to the TOC1 protein timeseries (Figure 7b). Most strikingly, their mRNA levels rise only after TOC1 levels have fallen substantially in the dark night of an LD cycle. Under LL, however, the *LHY* and *CCA1* mRNA levels rise from trough to peak while TOC1 protein is at peak levels, whereas the model shows simple, reciprocal regulation at this phase (Figure 7a), indicating that the bulk TOC1 protein measured by the reporter fusion does not match the TOC1 function represented in the full model (see Discussion).

**Figure 7.**
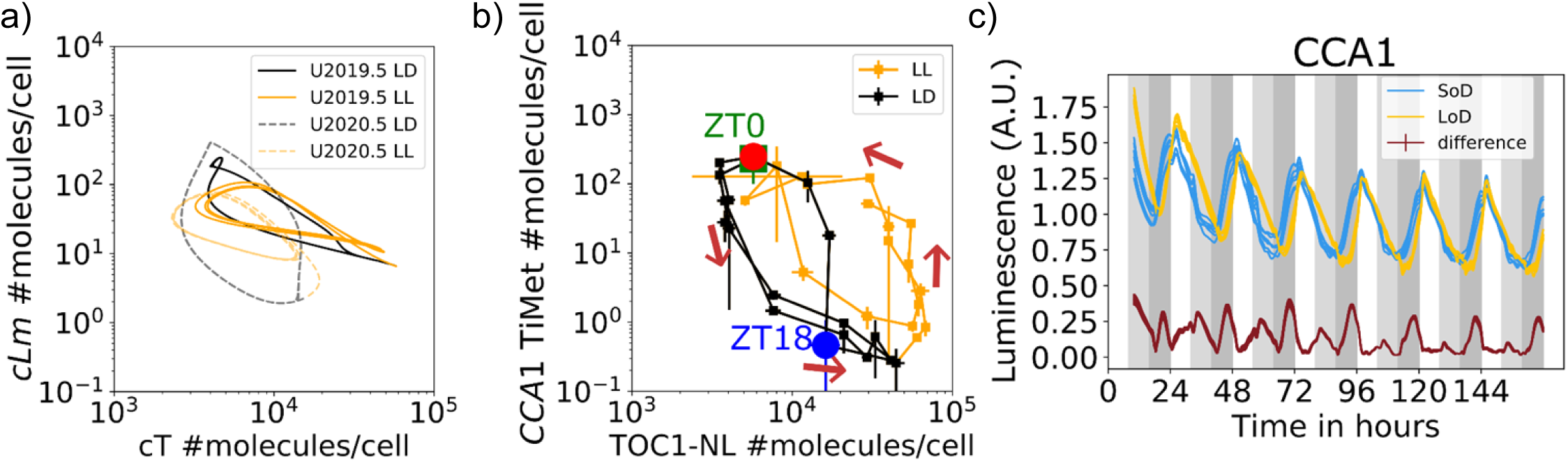
Dynamics of measured and modelled clock proteins. Phase plane diagrams show light-sensitive accumulation of TOC1 protein compared to its target *LHY* mRNA. (a) Variables *cT* and *cLm* in models U2019.5 (dashed lines) and U2020.5 (solid lines), and (b) TOC1 levels in extracts (as in Figure 6) and TiMet *LHY* mRNA data (Flis *et al*, 2015), under 12L:12D cycles (black lines) and constant light (yellow lines). Markers in (b) show the first (green) and second (red) ZT0 (lights-on) and the intervening ZT18 (mid-night) under 12L:12D, and the direction of time (arrows). The last data point in black is ZT12 under 12L:12D. In data (b), TOC1 levels then fall in 12L:12D but remain high under LL as *LHY* mRNA rises, which is not observed in the models (a). Error bars, 1 SEM. (c) *In vivo* recording reflects expected light-responsiveness, under short (8L:16D, cyan lines; SoD) compared to long photoperiods (16L:8D, yellow lines; LoD). Seedlings carrying the LHY protein reporter were grown for 10 of SoD or LoD in a multi-well plate then recorded hourly for 7 days in the same conditions, using an automated luminometer. Data were log-transformed and normalised to the mean of each timeseries (giving arbitrary units, A.U.). Data points for each seedling were connected with a cubic spline interpolation to facilitate comparision despite slight differences in sampling times. The absolute difference between the means (red line) emphasises the earlier rise of expression in the night under 8L:16D. White background, light interval; light grey, dark interval in 8L:16D only; dark grey, dark in both conditions.

### Protein timeseries data from *in vivo* imaging

ELF3 reporter lines were also created as outlined above and tested *in vivo* in seedlings, along with the four clock reporter fusions lines described above (Figure 6b, Supplementary Figure 6), and LUX (Supplementary Figure 4f). The *in vivo* luminescence levels from the LHY, CCA1, PRR7 and TOC1 reporters peaked within a 5-fold range, ranked LHY∼CCA1 > PRR7 > TOC1, consistent with the peak signals in extracts (Table 1). The bioluminescence signal *in vivo* peaked in phase with the timeseries in extracts for LHY and CCA1, or 2h later for PRR7 at ZT12 rather than ZT10 *in vitro*. The *in vivo* assay showed rapid falls of 10-30% in PRR7 and TOC1 fusion signals immediately after dusk (Figure 6e, Supplementary Figure 6). These were not evident in the *in vitro* timeseries, possibly due to its lower time resolution (Figure 6a). The *in vivo* assays of the four, well-expressed clock reporter genes showed quantitative consistency, despite some finer features that remain to be characterised, suggesting that the ELF3 reporter could also be characterised *in vivo*.

The transgenic ELF3 fusion plants expressed the reporter at similar or higher peak signal levels to the other four clock proteins, with a broad peak starting at ZT12 under constant light (Figure 6b), as expected from the RNA timeseries. The rhythmic amplitude was smaller than for the other clock protein fusions, under 2-fold, consistent with the simple model’s prediction and its underlying RNA timeseries data (Figure 2). These consistent features suggested that the peak ELF3 protein levels were also ∼40,000 molecules per cell (Table 1), by comparison to *in vivo* results for the other four clock proteins. Under LD cycles, the broad peak of ELF3 reporter signal was interrupted by an abrupt, transient signal decline of around 30% from ZT12 and by a minor peak at ZT2. The detailed dynamics of this reporter system need further characterisation, before such short-term features can be interpreted. For LUX, the transgenic line tested *in vivo* showed clearly-rhythmic expression with a peak at ZT12 as expected (Supplementary Figure 4f). The original line’s weak activity *in vitro* was qualitatively consistent with this rhythmic waveform (Supplementary Figure 5c). Calibrating the *in vivo* data as above for ELF3 suggested a peak of 5,000 LUX proteins per cell, a preliminary estimate far below the ∼100,000 predicted by the simple model (Table 1) and which awaits confirmation.

The *in vivo* assay allows rapid testing of multiple environmental conditions, illustrated here by repeating the assays under short- and long-photoperiod conditions, 8L:16D and 16L:8D (Figure 7c, Supplementary Figure 8). The phase of the night-time rise in LHY and CCA1 signal was clearly sensitive to the time of dusk, being delayed under long compared to short photoperiods, as was the waveform of TOC1, as expected (Edwards *et al*, 2010; Flis *et al*, 2016). A similar delay in the PRR7 phase was also expected (Flis *et al*, 2016) though photoperiod-responsive changes in the PRR7 reporter profile were less clear (Supplementary Figure 8). Overall, the NanoLUC reporter fusions can report temporal protein profiles *in vivo* and have potential to indicate absolute protein levels in some cases, by comparison to lines that have been tested by calibrated assays *in vitro*.

## Discussion

Converting mathematical models of the plant clock from arbitrary to absolute mass units is intended to improve model quality, by harnessing biochemical data (Urquiza-García & Millar, 2021). Absolute units are also helpful for any circuit engineering that includes previously-characterised components. The U2019.5 and U2020.5 models continue this process, by rescaling the previous model versions to match the clock protein levels, which we predicted from absolute RNA levels using the simple model. Protein levels measured in extracts of transgenic plants by the calibrated, NanoLUC reporter fusion technique were consistent with these predictions for LHY, CCA1, PRR7 and TOC1 (Figure 6, Table 1). The models’ simulated *K_d_*’s for DNA binding matched calibrated, empirical data from the literature for LUX binding within the EC but not for LHY/CCA1 (Figure 4, Table 2, Supplementary Table 4). The biochemical data that were available in absolute units, or which we could calibrate, were temporarily limiting, so we share the data and models (see Data Availability) for other researchers to pursue the remaining discrepancies such as those outlined below and in Supplementary Table 5.

### Absolute numbers of clock proteins

The success of the simple model suggests that its assumptions, together with the RNA data in absolute units and estimated protein degradation rates, were sufficient to estimate the dynamic, clock protein levels. In other words, the bulk levels of these clock proteins might be rather simply regulated, so the simple model’s approach could justifiably be repeated to estimate the levels of other proteins. Acquiring the necessary RNA data from absolutely-calibrated assays (Flis *et al*, 2015) could be easier than developing a calibrated, NanoLUC fusion reporter assay in extracts of transgenic plants (Urquiza-García & Millar, 2019).

The NanoLUC reporter assays in plant extracts estimated clock protein numbers that vary rhythmically from ∼1,000 to ∼100,000 copies per cell, for LHY, CCA1 and PRR7, with about half that peak level of TOC1. Individual plant cells probably have higher-amplitude protein cycles than these bulk figures. *in vivo* assays suggested that ELF3 and LUX levels peaked at 40,000 and 5,000 protein copies per cell respectively, though these preliminary estimates await confirmation. Our quantification of plant transcription factor abundance seems compatible with an estimate of 30,000 monomers of phytochrome B, which interacts with transcription factors (∼3000 phyB in each of up to 10 photobodies per nucleus, Kim *et al*, 2023), or ∼1 million copies of one of three histone H1 variants (AT2G30620; Probst *et al*, 2020) out of 160 million proteins in total per mesophyll nucleus (Heinemann *et al*, 2020). The equivalent ranges for mammalian, fungal and cyanobacterial clock proteins respectively include 1000-15,000 copies of PER2 (Narumi *et al*, 2016; Smyllie *et al*, 2016), low 10’s of copies of FRQ (Merrow *et al*, 1997) and 10,000-25,000 copies of KaiB (Chew *et al*, 2018; Kitayama *et al*, 2003) per cell or per nucleus. The apparently larger numbers of plant clock proteins might reflect functional requirements other than circadian timing, for example due to the cell biology of plant nuclei, or of these particular proteins and genomic target sites. Copy number data for other plant transcription factors are now required, to determine whether these clock proteins are unusual.

The protein numbers are larger than the ‘system sizes’ inferred only from the observed variability of Arabidopsis rhythms using stochastic models (Guerriero *et al*, 2012; Gould *et al*, 2018; Greenwood *et al*, 2022), similar to the equivalent comparison for the mammalian clock proteins (Forger & Peskin, 2005; Leise *et al*, 2012). The large protein numbers and high rhythmic amplitudes are expected to reduce the stochasticity of circadian timing at the cellular level but might limit its resetting by input signals (Pittendrigh *et al*, 1991), and contribute to the observed reproducibility of clock gene expression waveforms in plants under laboratory (Flis *et al*, 2015) and natural conditions (Nagano *et al*, 2012).

### Refining the modelled protein profiles

Our reporter fusion data were not used in model construction. The U2019.5 and U2020.5 model simulations match the timing of these molecular waveforms relatively well, reflecting the fact that the NanoLUC reporter data are similar to past expression studies in arbitrary mass units, which had informed model construction. Three examples illustrate the detailed questions on absolute protein levels, before we address the challenges of DNA binding (see Future Perspectives: investigating *K_d_ in vivo*).

First, our data allow a new focus on the amplitude of gene expression rhythms, which is functionally important for higher-order circadian properties. The 1000-fold amplitude of *CCA1* and *LHY* RNA expression waveforms under LD cycles (Flis *et al*, 2015) drives high-amplitude protein rhythms in the simple model but the full models have lower-amplitude protein rhythms, for example 3-fold to 8-fold lower for CCA1 and LHY (Table 1). Two technical factors likely contribute to this difference. The method for testing model parameters does not match the minimum and maximum points in particular, unless specific steps are taken (Troein *et al*, 2011; Fogelmark & Troein, 2014), in part because focussing on individual data points risks over-fitting. Our models also include some conservative choices that affect amplitude. Their clock genes respond to transcriptional regulators with moderate sensitivity (Hill factors fixed at 2), for example, whereas greater ultra-sensitivity can be observed in other systems. These choices might be relaxed based upon future evidence. Amplitude changes can also be informative, for example comparing the amplitude under LD cycles, where direct light/dark signals contribute, with only endogenous, circadian regulation under constant light (Figure 6A, Table 1). Balancing the contributions of many, parallel regulators is a common challenge in modelling complicated systems. However, the direction of amplitude change is simulated correctly by U2019.5 and U2020.5 for LHY, CCA1 and PRR7 (falling in LL) and the remaining absolute discrepancies are small, so validation studies with this level of accuracy might prove laborious.

Second, for TOC1, the full models closely match its protein rhythms and correctly increase in amplitude in LL relative to LD conditions (Figure 6a, Table 1), yet the model’s protein levels are not necessarily comparable with the fusion protein data. The phase plane diagrams (Figure 7b, Supplementary Figure 7) show *LHY* and *CCA1* mRNA levels in LL rise from trough to peak while TOC1 fusion protein levels are at or close to their peak, implying that measured TOC1 is not always an effective repressor of *LHY* and *CCA1* transcription. In the models, levels of these transcripts rise only as TOC1 protein falls (Figure 7a). The models represent only active TOC1 repressor, because there was no data to inform modelling of inactive TOC1 protein. The phosphorylation of TOC1 and other PRR proteins (Wang *et al*, 2010), in particular by CK1 (Uehara *et al*, 2019), might be sufficient to inactivate them at the end of the subjective night, though additional clock components could also contribute. Such phosphorylation might reflect the “phospho-dawn” observed in proteome-wide, rhythmic protein phosphorylation in the green lineage (Noordally *et al*, 2018; Kay *et al*, 2021; Krahmer *et al*, 2022; Noordally *et al*, 2023).

The full models depart furthest from the data in the third example, of ELF3 under LD cycles. *ELF3* RNA amplitude is only around 10-fold (Flis *et al*, 2015), or 1.7-fold for the ELF3 protein reporter *in vivo* under LD cycles (Figure 6b), identical with the amplitude predicted from the *ELF3* RNA rhythm by the simple model (Table 1). Other assays also detected significant ELF3 protein at all phases in Arabidopsis, consistent with a modest amplitude (Hicks *et al*, 2001; Nieto *et al*, 2022). However, the full models simulated 7- or 20-fold ELF3 protein amplitudes in LD, with 10-fold less protein at the trough than predicted from the RNA (Figure 3). The full models regulate ELF3 using the observed light regulation of ELF3 stability by the COP1 system (Yu *et al*, 2008), which the models represent in detail (Pokhilko *et al*, 2011). ELF3’s partner proteins (GI, ELF4 and LUX) are also multiply regulated by light inputs, potentially contributing regulation in the plant that the current models partially ascribe to ELF3. This reinforces the need to quantify the protein partners and their various complexes (Pokhilko *et al*, 2011), for example by absolutely-calibrated mass spectrometry (Narumi *et al*, 2016), in timeseries (Krahmer *et al*, 2019), in order to constrain these, parallel regulatory mechanisms of the Evening Complex.

### Future perspectives: interpreting NanoLUC profiles *in vivo*

The NanoLUC fusion reporters promise a rich source of quantitative, protein-level timeseries data for the modelling of plant clock gene circuits, for example to understand the interaction of light-regulated translation with rhythmic RNA abundance (Piques *et al*, 2009; Seaton *et al*, 2018; Bonnot & Nagel, 2021), as well as the genetic analysis of other regulatory systems. The impact of cellular factors on bioluminescence *in vivo*, such as variation in the substrate O_2_ concentration in transgenic plants, remains to be established in detail. Sensitivity to such factors in principle offers the opportunity to report other cellular parameters (Aflalo, 1991; Feord *et al*, 2019) but also affects the detailed interpretation of *in vivo* imaging data, such as the rapid fall in fusion reporter signals upon transition to darkness (Figures 6b, Supplementary Figures 6, 8). Studies using nominally-constitutive expression constructs will be required to interpret these details of the waveform, as previously for the firefly luciferase reporter (Millar *et al*, 1992; Van Leeuwen *et al*, 2000; Edwards *et al*, 2010).

### Future perspectives: investigating *K_d_ in vivo*

Tackling the model recalibration as a “Fermi problem” (Phillips & Milo, 2009) depends upon many published studies, and guarantees that many refinements to our approach will be possible (Supplementary Table 5). The full models’ simulated *K_d_* values for the Evening Complex binding to its target genes (0.3-8.6 nM) overlapped the calibrated range of *K_d_*’s for the LUX DNA-binding domain in gel-shift assays (1.3-3.2 nM) (Figure 4, Table 2,, Supplementary Table 4). No closer agreement could be expected given the approximations involved. This result depends on LUX levels and on the fraction of LUX protein that contributes to bind DNA in the EC, which emerges in the models only from indirect, functional constraints. For example, the EC must act relatively slowly in its negative feedback on the *LUX* and *ELF4* promoters (modelled by slow and incomplete LUX incorporation), in order to set the 17h period of rhythms in *lhy;cca1* double mutant plants (Locke *et al*, 2005; Pokhilko *et al*, 2012; Urquiza-García & Millar, 2021). The equivalent incorporation of some mammalian clock proteins, in contrast, has been directly tested (Aryal *et al*, 2017; Koch *et al*, 2022).

Our experimental data supported the simple model’s estimates of clock protein levels for LHY and CCA1 (Table 1), which bind DNA directly in the full model. These protein levels simulated *K_d_* values that were 1000-fold higher (lower binding affinity) than the measured *K_d_*values *in vitro*, extrapolated to the observed CCA1-binding regions (Figure 4, Table 2). Among many possible contributions to this discrepancy, the *in vitro* data tested single binding sequences, whereas the total protein number in cells has evolved with many specific binding sites and non-specific binding, titrating the available protein across the entire genome. The number of specific binding sites measured by ChIP-seq is close to 1000 (Adams *et al*, 2018; Ezer *et al*, 2017; Kamioka *et al*, 2016; Nagel *et al*, 2015). The mathematical form of our clock models assumes that the clock protein pools are large enough that the protein consumed by binding to target sites has negligible effect on the available protein pools. That assumption could have been contradicted if just a few thousand clock proteins were present per cell, close to the number of binding sites. In that counterfactual case, a higher binding affinity (lower *K_d_* value) would be required in our models to recognise that additional sites were competing to bind the clock protein, though these sites were not explicitly represented in the model. Recalibrating the model’s *K_d_* values using the observed number of CCA1 binding sites gave affinity values that overlapped with the *in vitro* estimates for binding to the same ChIP-seq regions (Figure 4, Table 2), another remarkable result. However, the simple model and the experimental measures in fact agreed on protein numbers around 100,000 per cell. Sequestering of clock proteins at around a thousand observed ChIP-seq sites is not expected to alter these protein concentrations significantly, consistent with our current models’ assumptions. It is possible that the measured, bulk clock protein levels might over-estimate the protein available for promoter binding, due to protein partitioning outside the nucleus (Yakir *et al*, 2009), clustering of proteins within the nucleus or a processing step akin to the formation of a small EC pool from a fraction of the bulk LUX protein. Otherwise, the high DNA-binding affinities of CCA1 and LHY measured *in vitro* contrast with the lower affinities (higher *K_d_* values) required in the mechanistic models.

The dissociation constant *K_d_* is a ratio of the binding and dissociation rates, *K_on_* and *K_off_*, either of which could be altered in the relevant *in vivo* conditions compared to the *in vitro* assays. The binding rate could be reduced if the accessibility of promoter regions is significantly lower *in vivo,* where nucleosomes and other chromatin or regulatory proteins can limit binding rate compared to naked DNA *in vitro*. ChIP-seq from intact chromatin (Adams *et al*, 2018) revealed many fewer LHY binding sites (around 700 sites) than the binding potential of LHY from DAP-seq on naked genomic DNA (around 18,000 sites)(O’Malley *et al*, 2016), for example. The dissociation rate could also be increased *in vivo* if the minority of bound proteins are removed from the promoter by nucleosomes or by an active degradation process (Spoel *et al*, 2009). Quantifying these biochemical mechanisms *in planta* is technically challenging. Our models should help to understand their effects at larger scales.

### Future perspectives: models informed by genome sequence

The absence of measured *K_d_*s for clock protein binding to target promoter sequences led us to estimate these affinities from protein-binding microarray data (Figure 5), an approach that should be applicable to other transcription factors, pending direct assays of binding *in vivo*. Our results analyse the CCA1 and EC binding sequences identified by ChIP in any of the model’s target genes (Table 2, Supplementary Table 4) but the approach applies equally to any sequence variant. Figure 8a shows the predicted effect of observed, natural variation in the CCA1-binding sequences in the promoter region of *GIGANTEA (GI)* among Arabidopsis accessions sequenced by the 1001 Genomes Project (Weigel & Mott, 2009). Any sequence-driven analysis that informs a parameter in the full model is within the scope of this approach, so a growing number of genome sequence variants might become interpretable. Alternative RNA splicing is a sequence-dependent regulatory mechanism (James *et al*, 2018), for example, which underlies the control of LHY translation that forms part of the plant clock’s response to temperature (Gould *et al*, 2013). For Arabidopsis, it seems premature to expect that DNA-binding affinities might be predictable from sequence variants in the DNA-binding domains of clock proteins, informed by biophysical data and sequence-base protein structure predictions (Jumper *et al*, 2021), but this might be tractable in other systems (Narasimamurthy *et al*, 2018).

**Figure 8.**
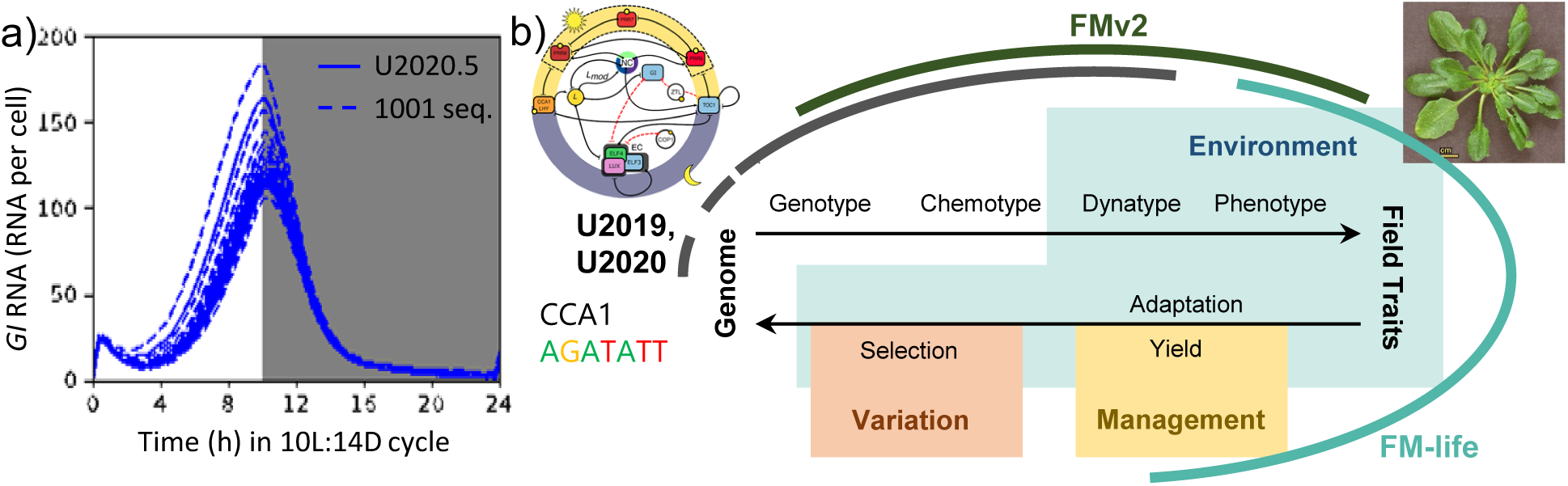
Connecting genome sequence variation to phenotypic effects across scales. Natural genetic variation in promoter sequences is predicted to alter the molecular phenotype (dynatype). (a) *GI* transcript levels under 10L:14D cycles were simulated in U2020.5 (white region, light interval; shaded region, dark interval). The *Kd* for CCA1 binding to the promoter of *GI* was calculated using the EMA matrix (Figure 5), for all *GI* promoter sequences from the 1001 Genomes Project (dashed lines). The model retained the default *GI* gene (solid line, second highest peak), while a second copy that simulated only *GI* RNA production was tested with the *Kd* for each promoter sequence and plotted (dashed lines). The range of dynamics shown reflects only altered *GI* transcription rates, without the effects of altered GI protein dynamics. (b) Connecting models (adapted from Millar, 2016 Figure 4c). The central circuit represents the conceptual steps as a genome, in cells, builds organismal traits (upper arrow) in a given environment (green shading). Those traits and potentially management inputs (yellow), in populations, lead to selection on genome sequences (lower arrow). The black, green and cyan arcs represent, respectively, the clock gene circuit models such as U2020 (schema from Urquiza-García & Millar, 2021 Supp Fig 3b), the Framework Model version 2 for simple environments (FMv2; Chew *et al*, 2022), and the FM-life model for recorded, natural environments (Zardilis *et al*, 2019). This paper illustrates how a clock gene circuit model can incorporate genome sequence data (dashed black arc), *via* promoter sequences that alter CCA1 binding, as in (a). Others might connect such models in future and add simulated genetic variation (pink), in order to explain and predict both the operation and evolution of the plant clock.

Sequence-informed, mechanistic models such as U2019.5 and U2020.5 promise to link these data on genetic biodiversity quantitatively to their effects at larger scales. *GI* expression both contributes to the photoperiodic control of flowering among *A. thaliana* accessions (de Montaigu *et al*, 2015) and has known and modelled molecular mechanisms (Seaton *et al*, 2015). The waveforms of Arabidopsis clock gene expression are already linked to whole-organism traits in the Framework Models (Kinmonth-Schultz *et al*, 2019; Chew *et al*, 2022). An initial extension to ecological data shows the potential to understand evolutionary adaptation in mechanistic terms (Burghardt *et al*, 2015; Zardilis *et al*, 2019). In the context of the Framework Models (Figure 8b) such as our FMv2 and FM-Life, a sequence-informed clock model might predict the change in flowering time in particular environmental conditions, and hence the phenology of flowering in nature. If population genetics (genetic variation and selection) can be incorporated into this system of models in the future, then plant science might make sense of the plant circadian system quantitatively and mechanistically, in the light of evolution (Dobzhansky, 1973; Millar, 2016).

## Materials and Methods

### Computational procedures

Analysis of published data to estimate translation and protein degradation rates for the simple model and analysis of protein-binding microarray data using Error-Model Averaging are described in the Supplementary Information. All the mathematical models were written in the human-readable language Antinomy and transformed into SBML files using the python package Tellurium (Choi *et al*, 2018). Rescaling parameters were introduced into the models U2019.3 and U2020.3 (Urquiza-García & Millar, 2021) to match protein levels to the levels predicted by the simple model, creating U2019.4 and U2020.4, respectively (Figure 3). The scaling factors were estimated using the least-squares method implemented by the minimize function of the python package lmfit (Newville *et al*, 2023), with a custom cost function that generates numerical solutions of the model, and also plots simulation results, using the Tellurium software (Choi *et al*, 2018). Unless modelled otherwise, clock proteins were assumed to be nuclear and their concentrations were calculated using a published nuclear volume (Tirichine *et al*, 2009) from our previous parameter compendium (Millar *et al*, 2019). The modelling software system was run in a Docker container, described by the Docker file shared in the data compendium (see Data Availability).

### Cloning and prototyping

The Arabidopsis Col-0 TAIR10 genome assembly was used a reference genome. The genomic region of clock genes was amplified from Col-0 genomic DNA with primers described in Supplementary Table 7, apart from *ELF3*. The fragments were then recombined into pDONR221 using BP II clonase (Invitrogen) and sequence determined by sanger sequencing (Genepool, Edinburgh Genomics, Edinburgh, UK). The genomic regions were recombined into pGWB601:NanoLUC-3FLAG-10His using LR clonase, as described (Urquiza-García & Millar, 2019). *ELF3* was cloned in the same destination vector using Gibson cloning with primers described in Supplementary Table 7. The constructs were then transformed into *E. coli* DH5α and selected on Spectinomycin. The candidate plasmids were tested for NanoLUC activity and rhythmicity (Supplementary Figure 3) by transfecting protoplasts as described in Supplementary Information (Hansen & Ooijen, 2016; Urquiza-García & Millar, 2019).

### Plant transformation

The pGWB601:XpX-NL3F10H vectors carrying genomic regions of interest (X in the plasmid name) were transformed using liquid nitrogen into *Agrobacterium tumefaciens* ABI (kindly provided by Prof. Seth Davis, York). Arabidopsis plants were transformed by floral dipping (Wang, 2015). Primary transformants were selected for 3:1 segregation of BASTA resistant:sensitive progeny, to identify single-insertion lines. Single-insertion transgenic lines were analysed for phenotypic complementation in the T3 generation. For each construct, 15 or more homozygous lines with segregation consistent with single-insertion events were tested for a circadian period closest to the wild-type control and, for LUX and ELF3, also for hypocotyl elongation (Supplementary Information; Supplementary Figure 4).

### *In vivo* luciferase imaging

Period determination of transgenic lines was performed by *in vivo* luciferase reporter gene imaging, essentially as in Southern et al. (2006), see Supplementary Information. Period analysis with the FFT-NLLS algorithm in the public Biodare2 resource (Zielinski *et al*, 2014, 2022) used data starting after the first day in constant light conditions. Transgenic lines with the closest period and relative amplitude error relative to the corresponding WT control lines were retained for further analysis.

*In vivo* luminometry of NanoLUC reporters was similar to (Urquiza-García & Millar, 2019). 4 sterilised seed of each homozygous transgenic line were sown per well of a white, flat-bottomed 96-well plate that contained 150µl of solid agar media (see Supplementary Information). After one day at 4°C, plates were incubated for 10 days in 12L:12D conditions and treated with 50 µl of 1:50 of furimazine substrate solution (Promega, Southampton UK): 0.01% Triton X-100. The plate containing 8 biological replicates per construct was then assayed by luminometry every 30 or 60 minutes, in an automated TriStar2 S LB 942 luminometer (Berthold Technologies, Harpenden, UK) at 21°C, and exposed to monochromatic blue and red LED light with a total 50 µmol m^-2^s^-1^, 12L:12D photoperiod between readings for three days, followed by constant light. The falling trend in the signal of all reporters over several days in constant light is assumed to result from decay of the substrate furimazine. *In vivo* data from TOC1 and CCA1 reporter plants were scaled such that the means matched the cognate *in vitro* data. This scaling was repeated on detrended *in vivo* data for ELF3 and LUX reporters, to estimate their protein copy numbers.

### Calibration of the NanoLUC assay in plant extracts

Protein produced from the MBP-NanoLUC-3FLAG-10His construct in *E. coli* was purified as previously described (Urquiza-García & Millar, 2019) and quantified by a linear version of the Bradford protein assay (Ernst & Zor, 2010). Col-0 Arabidopsis plants were grown in 5 cm diameter tissue culture dishes, using media and growth conditions as above, for 10-14 days and collected in 0.1 gramme fresh weight (gFW) aliquots in 2ml microcentrifuge tubes (Safelock^®^, Eppendorf), along with two, 2mm stainless-steel grinding balls. Sufficient MBP-NanoLUC3F10H protein was added to the tissue samples to correspond to starting levels of 0, 1×10^2^, 1×10^3^, 1×10^4^, 1×10^5^, 1×10^6^ monomers cell^-1^, using the previously-measured average of 25 million cells/gFW of leaf tissue (Flis *et al*, 2015). The aliquots were then frozen in liquid nitrogen and extracted as detailed below, for NanoLUC assays of transgenic plant tissue. A calibration curve was generated on each 96-well plate measured, using 4 biological replicates per NanoLUC dilution. The data were ln-transformed, and a linear regression to these data yielded the calibration standard for each plate.

### Generation of NanoLUC timeseries from plant extracts

Seed of transgenic lines were sown on solid media as described above. Growth and harvesting conditions matched the TiMet protocol (Flis *et al*, 2015). After two weeks under a 12L:12D photoperiod of cool white fluorescent light at 21 °C, robust young plants were transferred to F2+S Levington compost (Frimley, UK) and grown until they were 21 days old. Leaf rosettes were pooled from 5 plants per biological replicate in 2 ml microcentrifuge tubes (Safelock^®^, Eppendorf) containing two, 2 mm stainless steel grinding balls and flash-frozen in liquid nitrogen. The sample tubes were maintained under liquid nitrogen, ground twice using a Tissuelyser (Qiagen Ltd., Manchester), and returned to liquid nitrogen before extraction. On ice, 150 µl of BSII buffer (100 mM sodium phosphate, pH 8.0, 150 mM NaCl, 5 mM EDTA, 5 mM EGTA, 0.1% Triton X-100, 1 mM PMSF, 1× protease inhibitor mixture (Roche, Basel, Switzerland) and 5 μM MG132 (Stem Cell Technologies, Cambridge, UK)) was added similar to (Huang *et al*, 2016) but without phosphatase inhibitors. The tubes were thoroughly vortex-mixed, returned to ice, while each tube’s mass was individually measured which then was adjusted with BSII buffer to reach a tissue concentration of 0.4 gFw ml^-1^ as recommended by the manufacturer for the protease inhibitors (Sigma-Aldrich). The extracts were clarified by centrifugation at 20,000xg for 10 min. 20 µl of plant extract per well was added to 96-well plates that were pre-loaded with 80 µl of BI assay buffer (50 mM NaH_2_PO_4_, 0.3 M NaCl, pH 8.0 NaOH adjusted). 100 µl of 1:50 Furimazine:NanoGlow (Promega, Southampton UK) were added using a multipipette. After 10 min incubation at 21°C in a temperature-controlled growth room, the bioluminescence was measured in a Tristar2 S LB 942 luminometer (Berthold Technologies, Harpenden UK) at 21°C with a signal integration time of 1.5s. The number of NanoLUC molecules cell^-1^ was calculated from the bioluminescence values, based the calibration standard (see above).

## Data Availability

The live, updatable resources from this study are public on FAIRDOMHub.org at https://fairdomhub.org/investigations/570, structured according to the standard ISA hierarchy, including the rescaled models U2019.5 and U2020.5 in multiple formats, along with the empirical data, construct maps, scripts, Docker container for the software environment and other resources.

Upon publication, the same resources will be shared as a static Snapshot, formatted as a Research Object on FAIRDOMHub.org (doi: 10.15490/FAIRDOMHUB.1.INVESTIGATION.??), on the Zenodo repository (doi: 10.5281/zenodo.??) and on Edinburgh DataShare (doi’s to be added in proof??). The data should be cited using the relevant DOI (or the FAIRDOMHub URL), for example as: Urquiza-Garcia, U., Molina, N., Halliday K. J. & Millar, A. J. (2024). Absolute units in the circadian clock models of Arabidopsis up to U2020.5 [Data set]. Zenodo. https://doi.org/10.5281/ZENODO.??

The rhythmic expression data and period analysis used to select complemented lines (Supplementary Figure 4a-c; Supplementary Table 8) are available from the BioDare2 repository for chronobiology data (Zielinski *et al*, 2022) in the following three experiments:

1. ID 10848; prr79 CCR2:LUC period complementation using PRR7NL constructs; permalink: https://biodare2.ed.ac.uk/experiment/10848
2. ID 11139; Complementation experiment lines CCA1NL, LHYNL, TOC1NL; permalink: https://biodare2.ed.ac.uk/experiment/11139
3. ID 11043; LUXp:LUX-NL3F10H; permalink: https://biodare2.ed.ac.uk/experiment/11043

Along with data for *in vivo* analysis (Figure 6, Supplementary Figure 6)

4. ID 11391; Plate reader experiment CCA1 TOC1 NanoLUC; permalink: https://biodare2.ed.ac.uk/experiment/11391

## Acknowledgements

For the purpose of open access, the author has applied a Creative Commons Attribution (CC BY) licence to any Author Accepted Manuscript version arising from this submission. For their expert assistance, we thank the plant growth facility, especially Sophie Haupt; Sarah Hodge with ultra-low-light imaging, Katalin Kis with laboratory methods and Dr. Marissa Valdivia Cabrera for curation of transgenic lines. For generously sharing materials, we thank Prof. Tsuyoshi Nakagawa (Shimane University, Japan) for providing pGWB vectors, Prof. Seth Davis (York University, UK) for the ABI strain and Prof. Norihito Nakamichi (Nagoya University, Japan) for sharing seeds of Col-0 CCA1-LUC lines.

## Author contributions

UUG, NM and AJM designed the study; UUG performed experiments; UUG performed modelling; UUG and AJM analysed experimental and theoretical results; UUG and AJM curated the data; UUG and AJM drafted the manuscript with support from all authors in review and editing; KJH and AJM supervised the work; UUG, KJH and AJM acquired funding.

## List of Supplementary Tables

Supplementary Table 1.

Estimated translation *s* and degradation *k* parameters for the simple protein model Supplementary Table 2

Parameter values of the U2019.5 model, and corresponding nuclear concentrations for *Kd* parameters *g1-g16*

Supplementary Table 3

Parameter values of the U2020.5 model, and corresponding nuclear concentrations for *Kd* parameters *g1-g16*

Supplementary Table 4

All DNA-binding dissociation constants (*K_d_*) from models U2019.5 and U2020.5 simulation and derived from experimental data using EMA.

Supplementary Table 5

Predictions, limitations, gaps and areas for future work.

Supplementary Table 6

Cloning of genomic fragments of clock genes for C-terminal fusions with NanoLUC-3FLAG-10His.

Supplementary Table 7

Primer sequences for genomic cloning of clock genes.

Supplementary Table 8

Circadian periods of host plants and complemented, transgenic lines carrying protein reporter fusions, measured using LUC+ reporter genes.

## Supplementary Information, contents

1. The simple model to predict clock protein levels from mRNA data
2. Deriving *K_d_*for promoter sequences by integrating Protein-Binding Microarray data and Surface Plasmon Resonance data.
3. Using energy matrices obtained by Error-Model Averaging
4. Assigning *K_d_* values to genes in the model
5. Construction of transgenic plants with NanoLUC-tagged clock proteins.

5.a Preliminary testing in protoplasts
5.b Mutant complementation
6. Supplementary Information References

## Figures and Supplementary Figures for

**Supplementary Figure 1.**
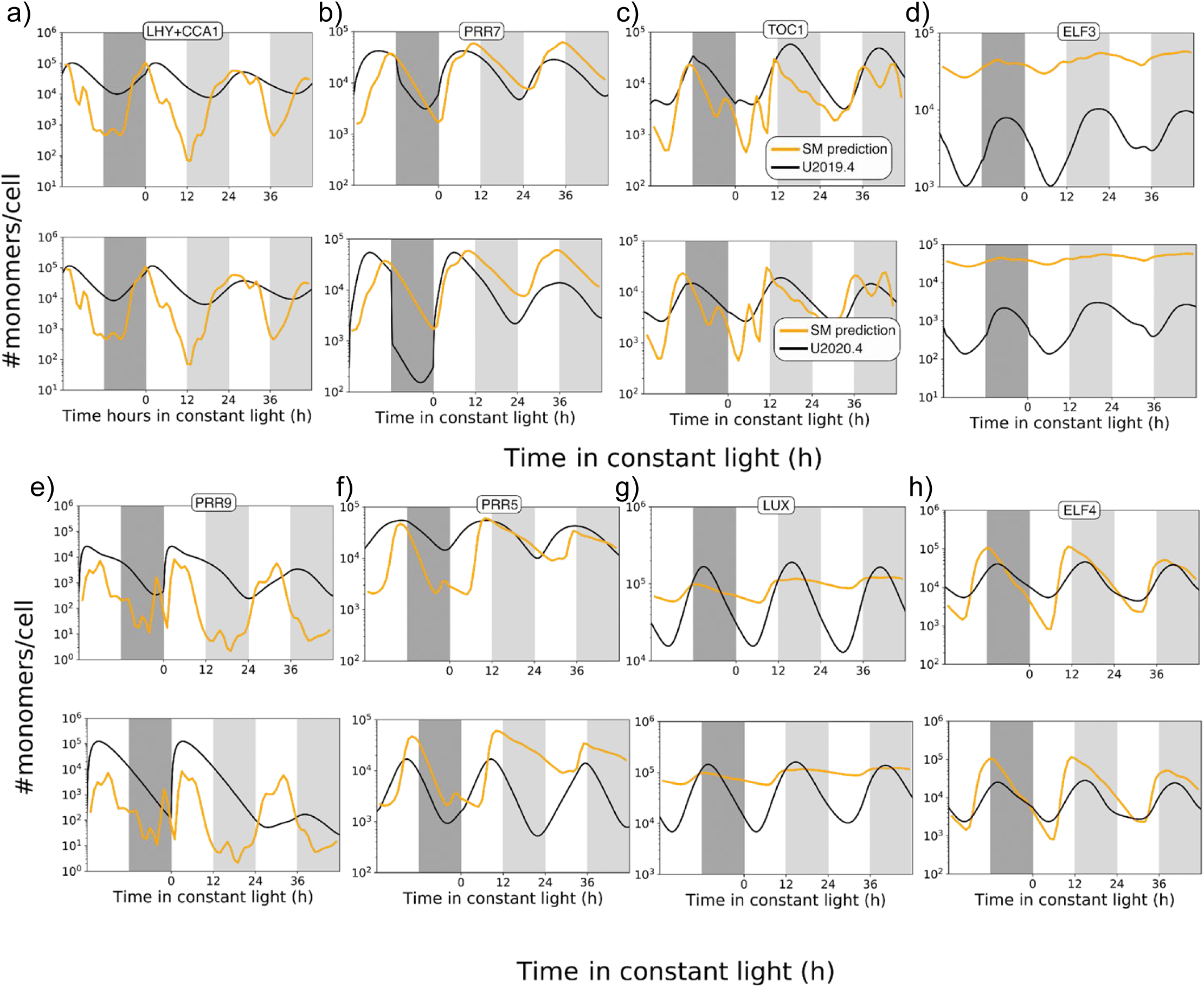
Rescaling protein levels in model U2020.4. Model simulations (black lines) were rescaled to fit the protein waveforms predicted from the simple model (SM, orange lines), for a) LHY/CCA1 (*cL*) fitted to the sum of predicted LHY + CCA1 protein, b) PRR7, c) TOC1, d) ELF3, e) PRR9, f) PRR5, g) LUX and h) ELF4. For each model component, the upper panel shows model U2019.4 (as in Figure 3), and the lower panel model U2020.4, simulation of protein dynamics in molecules per cell (black lines). The models were entrained to ten 12L:12D cycles prior to the interval plotted. Plots are scaled differently for each protein.

**Supplementary Figure 2.**
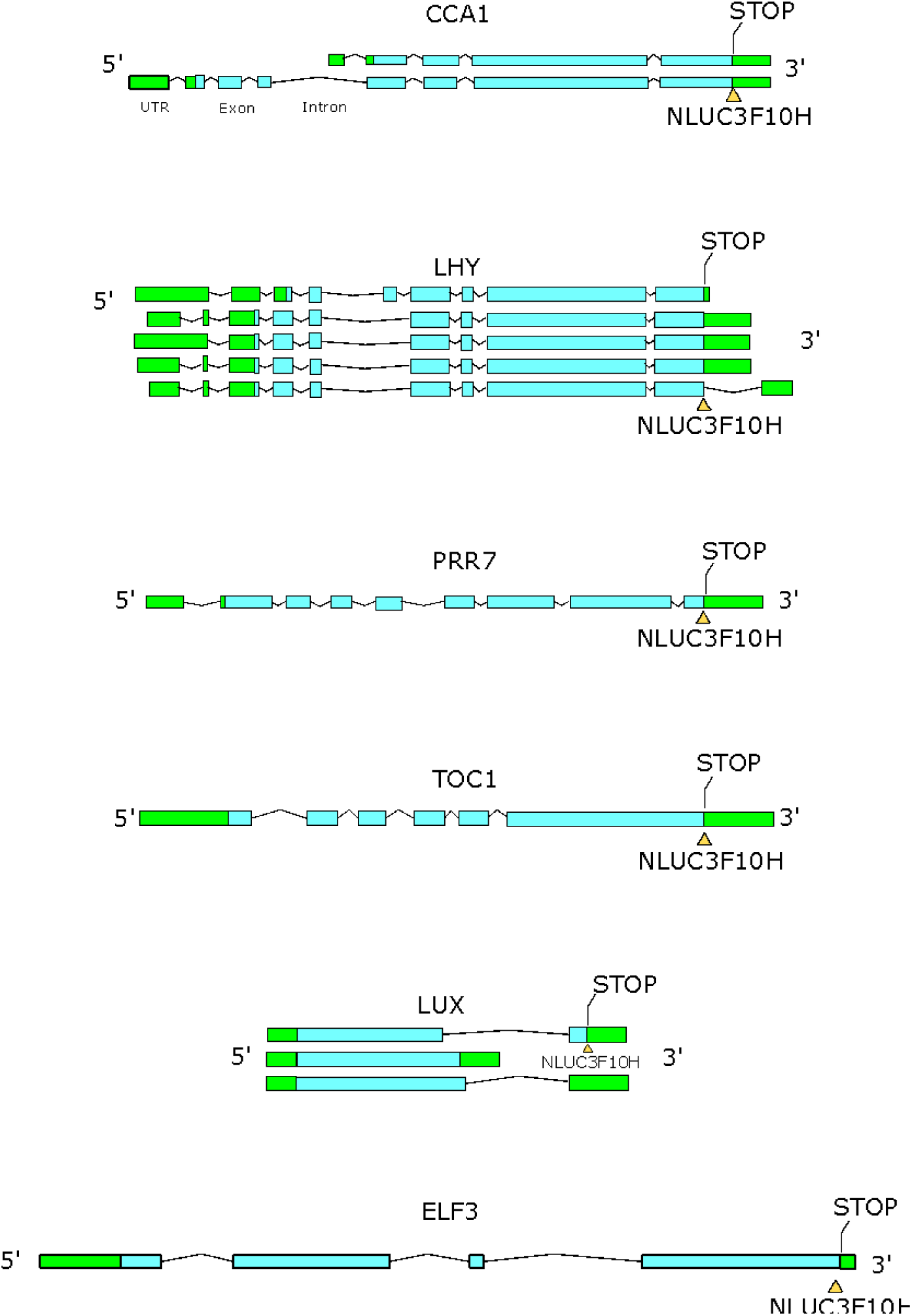
Reporter insertions into clock genomic regions. Genomic regions from Arabidopsis Col-0 were cloned in pDONR and subcloned into plasmid pGWB601::NanoLUC-(FLAG)_3_-(His)_10_::Tnos by means of Gateway cloning, as described (Urquiza-García & Millar, 2019). Genomic regions covering the most abundant RNA isoforms were selected for insertion of the NanoLUC-(FLAG)_3_-(His)_10_ protein (NLUC3F10H), with positions and primers listed in Supplementary Tables 6 and 7.

**Supplementary Figure 3.**
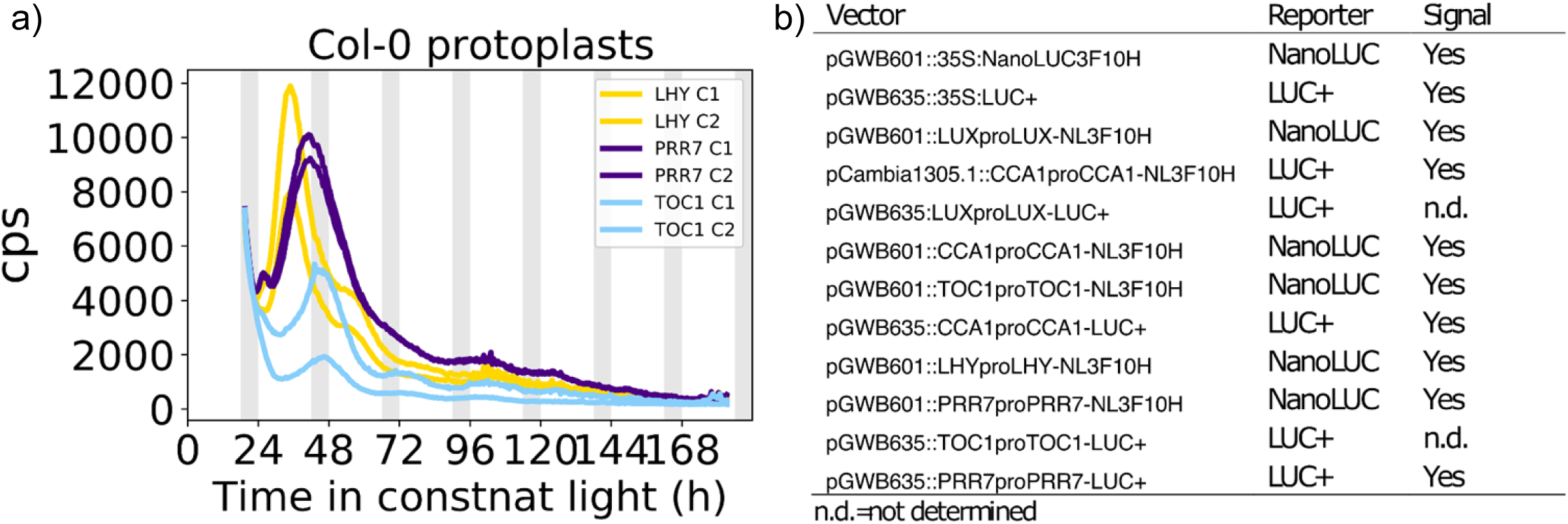
Rapid testing of protein fusion constructs in protoplasts. (a) Mesophyll protoplasts were extracted from 4 week-old plants of Arabidopsis thaliana Col-0 grown under 16L:8D conditions (Hansen & van Ooijen, 2016), transiently transformed with 10µg of each plasmid DNA and recorded under constant light in an automated luminometer, reading in counts per second (cps). Sample data are shown for NanoLUC fusions to LHY, PRR7 and TOC1. The first peaks (24-48h) show the expected sequence, LHY-PRR7-TOC1. Signals declined quickly but remained rhythmic. Shaded region, anticipated dark interval during plant growth. (b) All the reporter fusions tested showed rhythmic signals in this assay, though two LUC+ fusions were not tested (n.d.).

**Supplementary Figure 4.**
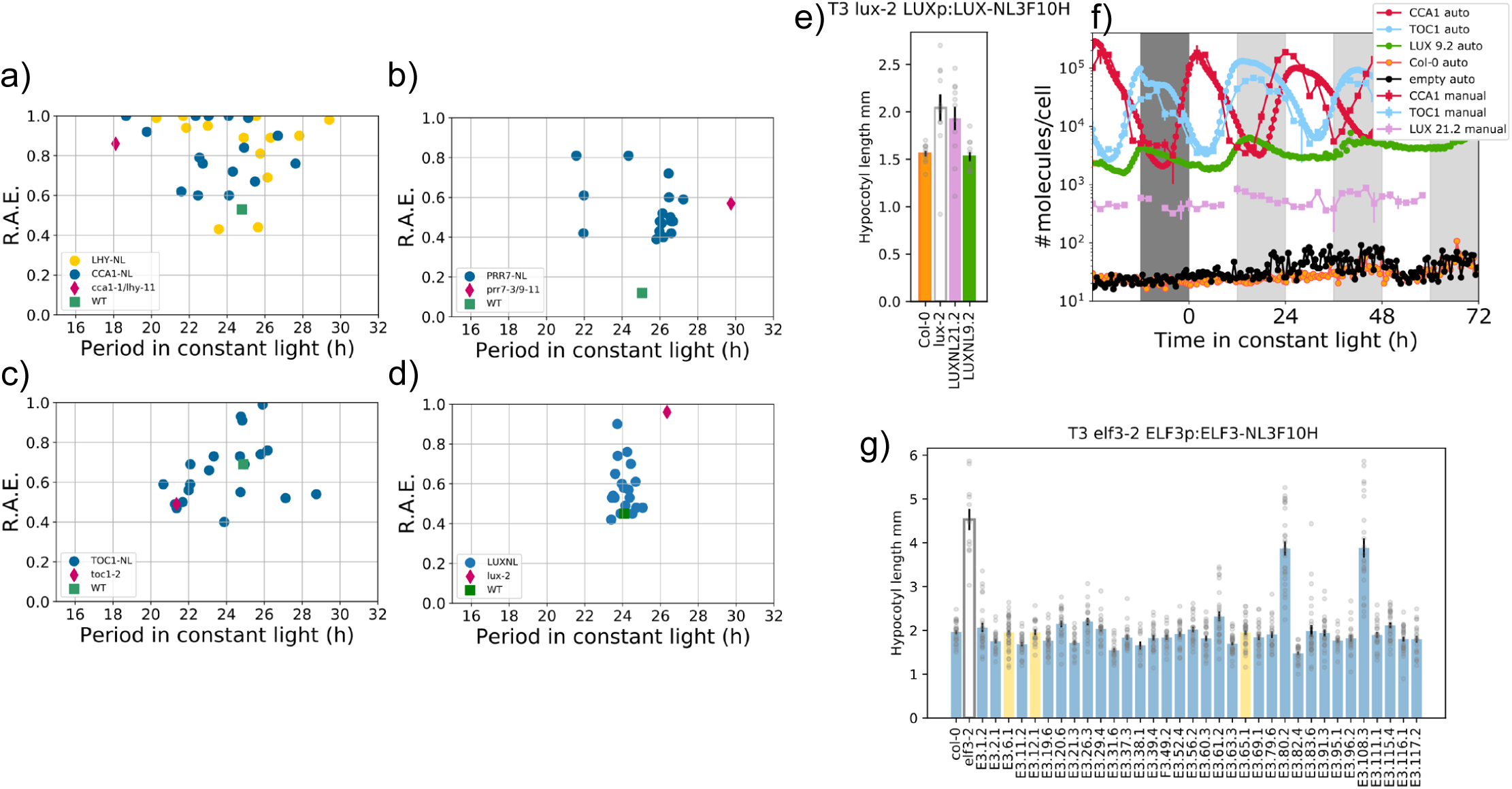
Reporter fusion constructs rescue clock mutant phenotypes. Each NanoLUC protein reporter line was selected for the rescue of a clock gene mutant phenotype. (a-d) Circadian rhythms in reporter lines were monitored by *in vivo* imaging of seedlings under constant light, using the firefly LUC transcriptional reporter included in the background of each mutant. Each data point represents the period and Relative Amplitude Error (R.A.E.) of a group of seedlings (n=10) from an independent, single-insert, homozygous line in the T3 generation for the constructs listed, compared to the mutant host (red diamond) and wild type (green square) controls. a) CCA1p:CCA1-NL3F10 (blue) and LHYp:LHY-NL3F10H (yellow) in the mutant background *cca1-1/lhy-11 CCA1p:LUC*. b) PRR7p:PRR7-NL3F10H in *prr7-3/prr9-11 CCR2:LUC*. c) TOC1p:TOC1-NL3F10H in *toc1-2 CCA1p:LUC*. d) LUX2p:LUX-NL3F10H in *lux-2 CAB2:LUC*. (e-g) In a second round of selection, (e) a further LUX protein reporter line was selected (line 9.2, green), which was complemented to the Col-0 hypocotyl length (orange) whereas the line 21.2 (pink) retained the long hypocotyl phenotype of the *lux-2* mutant parent (open bar). (f) *in vivo* data (‘auto’) from line 9.2 under LD and LL (green), after the same detrending and rescaling as for CCA1 (red) and TOC1 (blue)(see Methods), suggested an expression level about 7-fold higher than line 21.2 (pink) tested in extracts (‘manual’). Col-0 controls (orange) had the same low background signal as an empty well (black). (g) the ELF protein reporter lines (yellow) were selected based on the hypocotyl phenotype rescued to match Col-0 (far left, blue) and clearly rescue the long hypocotyl of the *elf3-2* mutant control (open bar).

**Supplementary Figure 5.**
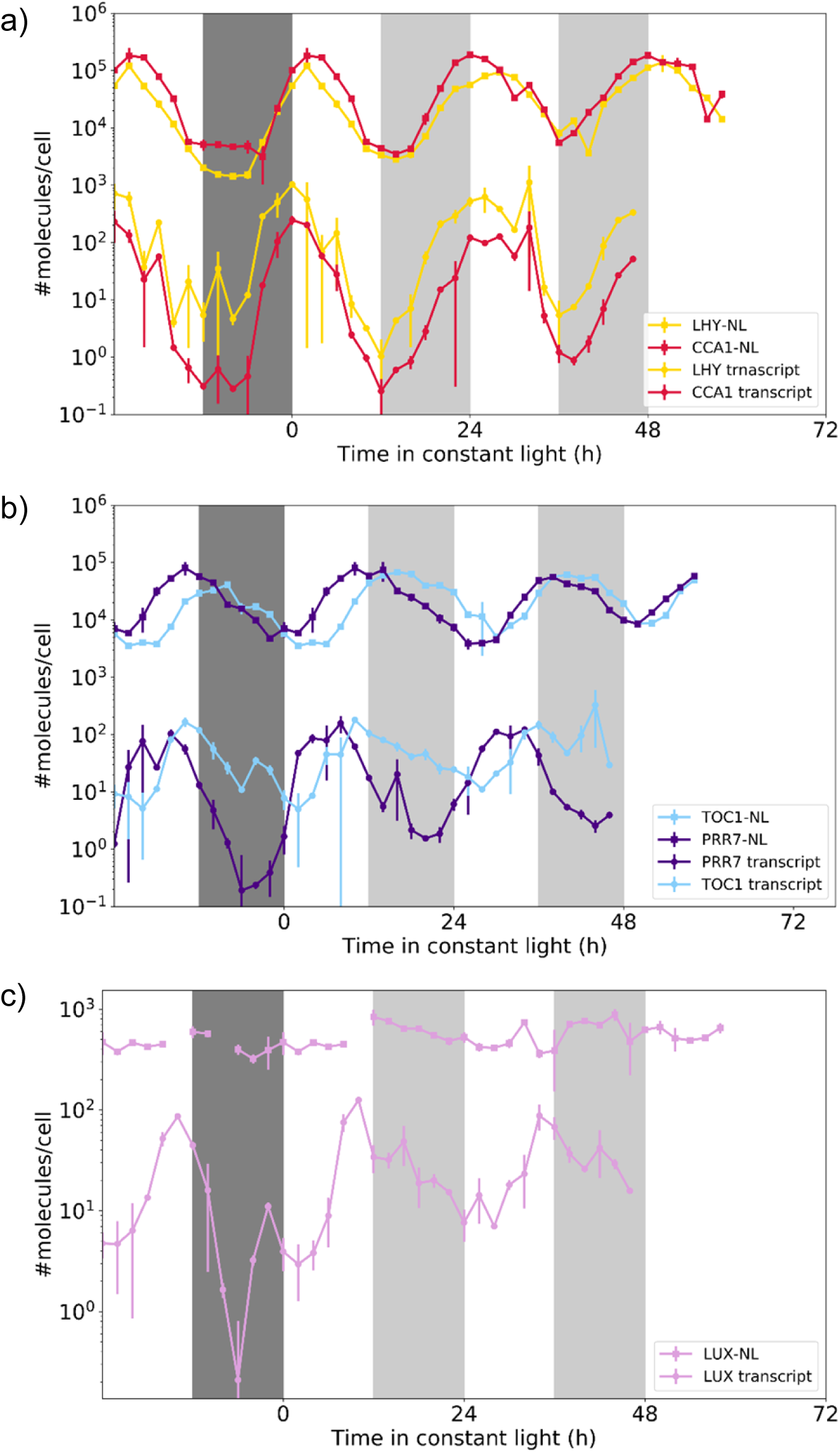
Timeseries of clock protein copy numbers relative to mRNA. Reporter protein levels in plant extracts (data as in Figure 6) were measured in calibrated NanoLUC assays for (a) LHY and CCA1, (b) PRR7 and TOC1 and (c) LUX line 21.2, under a 12L:12D cycle followed by constant light from time 0h. Note that this LUX line was only partially complemented, see Supplementary Information. Protein levels are compared to RNA levels in the TiMet data set, each in units of molecules per cell. Protein data are means of biological triplicates, RNA data are means of duplicates, error bar = 1 SEM. Light interval, white background; dark interval, dark grey shading; anticipated dark interval during constant light, light grey shading.

**Supplementary Figure 6.**
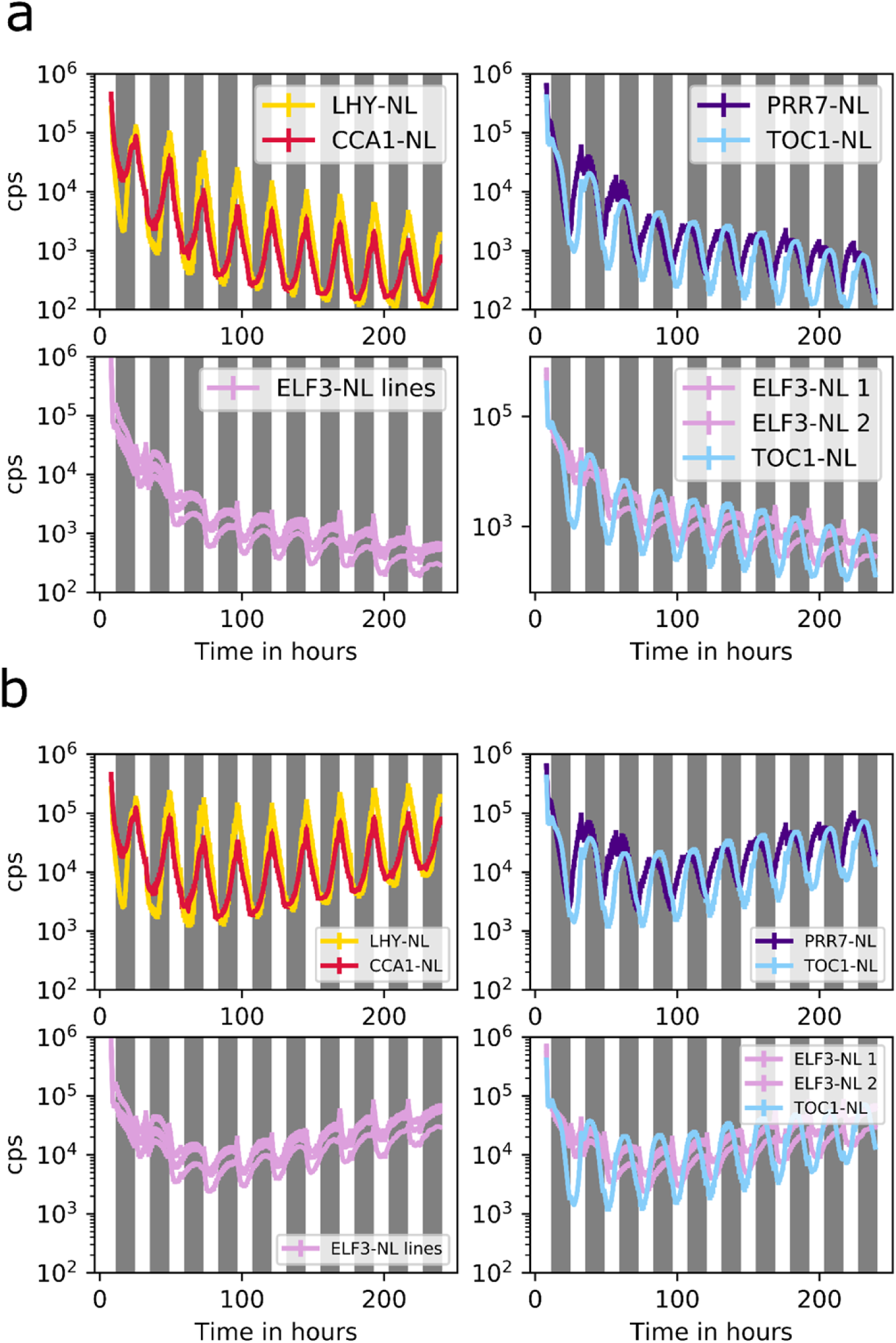
Long-term monitoring of protein fusions *in vivo*. (a) NanoLUC activity was measured (as in Figure 6 d-f) over ten 12L:12D cycles, in seedlings carrying the protein reporters indicated, for LHY (yellow), CCA1 (red), PRR7 (purple), TOC1 (cyan) and in three ELF3 reporter lines (pink), hourly in an automated luminometer. Each trace is from a micro-well plate seeded with 4 seeds per well and incubated under ten, 12L:12D cycles, treated with furimazine substrate and assayed for a further 10 cycles. (b) The falling trend due to furimazine decay was removed from the *in vivo* signals. A single exponential decay rate was estimated from a large data set of the CCA1 protein reporter and applied to detrend all the data sets. All the lines show acute responses to light-dark transitions.

**Supplementary Figure 7.**
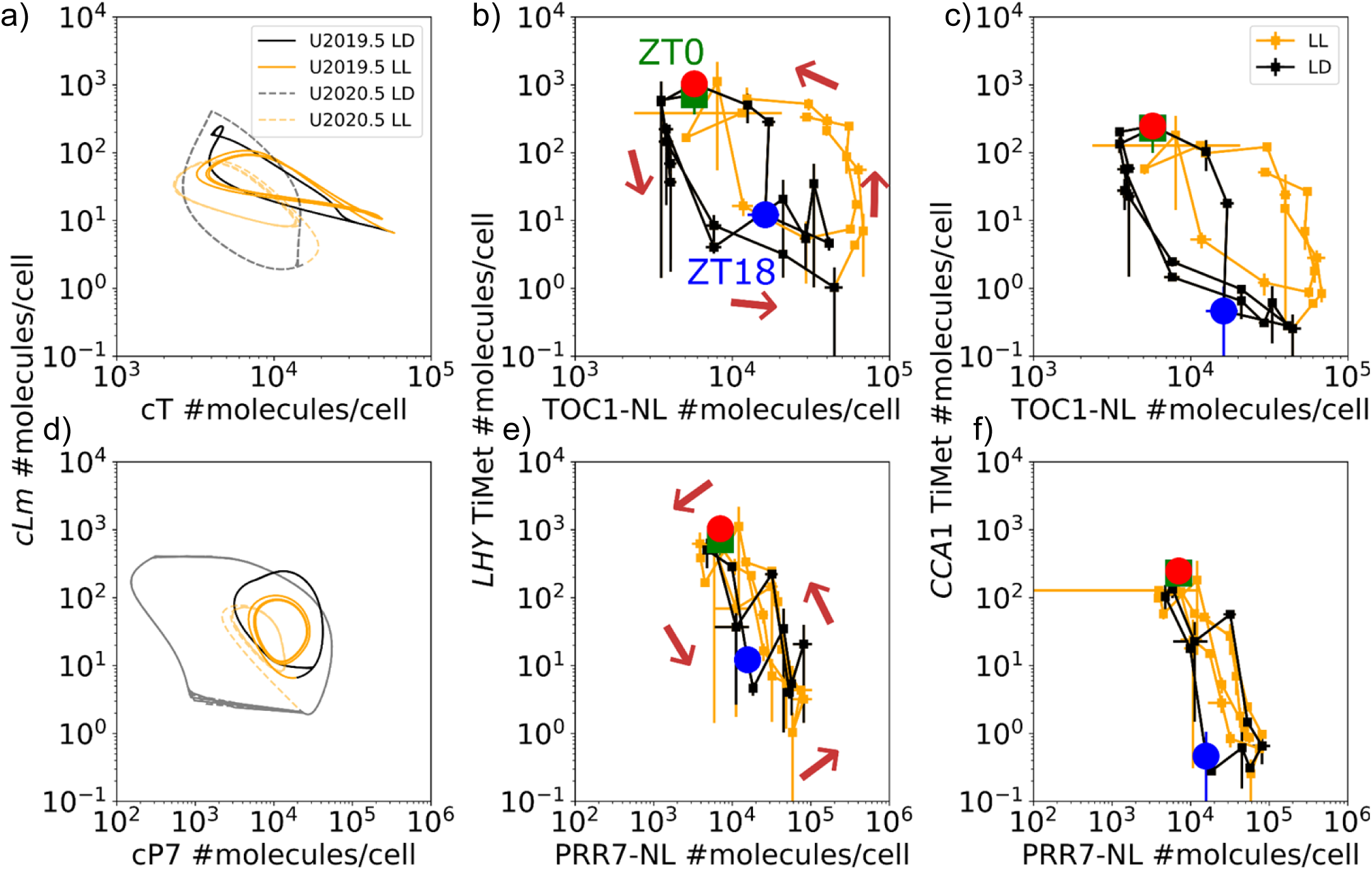
Regulation of *LHY* and *CCA1* by TOC1 and PRR7. Phase plane diagrams compare the accumulation of PRR transcriptional repressor proteins compared to their target mRNAs (as in Figure 7). (a) Variables *cT* and *cLm* in models U2019.5 (dashed lines) and U2020.5 (solid lines), (b) TOC1 levels in extracts (as in Figure 6) and TiMet *LHY* mRNA data, (c) TOC1 levels in extracts and TiMet *CCA1* mRNA data (Flis *et al*, 2015), under 12L:12D cycles (black lines) and constant light (yellow lines). TOC1 levels are lower when *CCA1* mRNA levels in 12L:12D than under LL (c), as for *LHY* mRNA levels (b). (d) Variables *cP7* and *cLm* in models U2019.5 (dashed lines) and U2020.5 (solid lines), (e) PRR7 levels in extracts (as in Figure 6) and TiMet *LHY* mRNA data, (f) PRR7 levels in extracts and TiMet *CCA1* mRNA data, under 12L:12D cycles (black lines) and constant light (yellow lines). These variables are anti-correlated in the data (e, f) but plot a more circular cycle orbit in the model simulations (d). The simulated orbits also contract under LL (d), whereas both mRNA and protein amplitudes are maintained in the data (e, f). Markers in (b) show the first (green) and second (red) ZT0 (lights-on) and the intervening ZT18 (mid-night) under 12L:12D, and the direction of time (arrows). The last data point in black is ZT12 under 12L:12D. Error bars, 1 SEM.

**Supplementary Figure 8.**
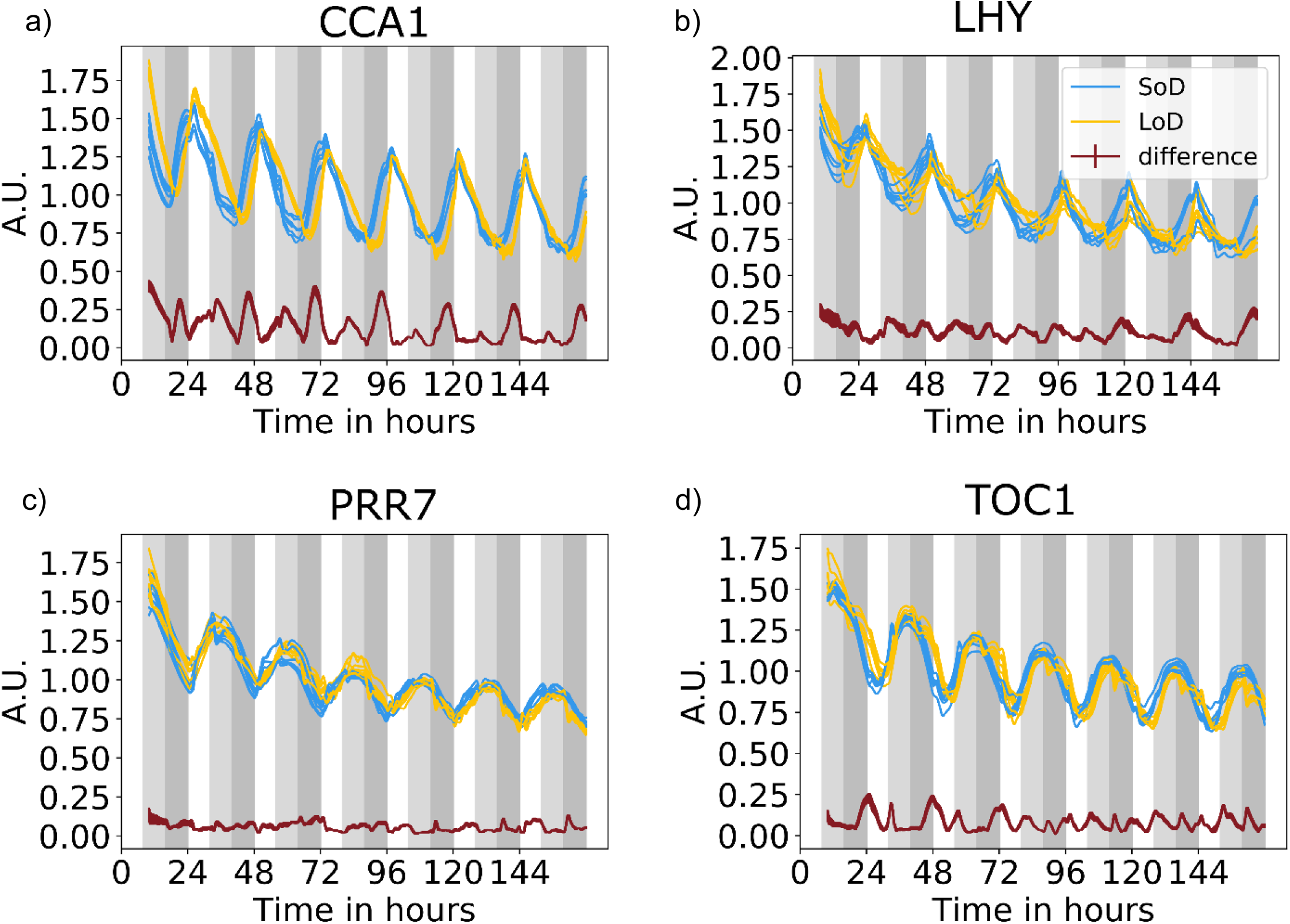
Clock protein dynamics respond to photoperiod *in vivo*. *In vivo* recording reflects expected light-responsiveness, under short (8L:16D, cyan lines; SoD) compared to long photoperiods (16L:8D, yellow lines; LoD), as in Figure 7c. Seedlings carrying reporter protein fusions to (a) CCA1, (b) LHY, (c) PRR7 and (d) TOC1 were grown for 10 of SoD or LoD in a multi-well plate and recorded hourly for 7 days in the same conditions, using an automated luminometer. Data were log-transformed and normalised to the mean of each timeseries (giving arbitrary units, A.U.). Data points for each well were connected with a cubic spline interpolation to facilitate comparison despite slight differences in sampling times. The absolute difference between the means (red line) shows the earlier rise of expression in the night under 8L:16D. White background, light interval; light grey, dark interval in 8L:16D only; dark grey, dark in both conditions.

