## Supplementary Information for "Abundant clock proteins point to missing molecular regulation in the plant circadian clock"

#### 1. The simple model to predict clock protein levels from mRNA data

Timeseries of mRNA transcript levels for *CCA1, LHY, PRR9, PRR7, PRR5, TOC1, ELF3, ELF4* and *LUX* were considered (Flis et al., 2015), in the form of a cubic-spline interpolation of ln-transformed transcript data. The cubic spline interpolates between experimental timepoints, while we have previously shown that the experimental errors within this dataset are close to normally distributed after log transformation (Urquiza-García & Millar, 2021). *GIGANTEA* data was not included because there was insufficient protein timeseries data in the literature to estimate the protein's degradation rate, in order to inform parameter *k* in the simple, data-driven model,

$$\frac{dP}{dt} = sM(t_i) - kP \quad (1)$$

Where $P$ stands for the amount of the protein of interest, $s$ is its translation rate, $M(t_i)$ its cognate mRNA transcript level in absolute units at time $t{=}i$ and $k$ is the decay constant of the protein. Translation rates follow from the number of codons (specific to each Open Reading Frame, ORF), the ribosomal density and ribosome elongation rate, each of which is a constant. We follow past work (Piques et al., 2009) in correcting a translation rate estimate for mammalian cells, as follows. The reference elongation rate at 25-26ºC for animal cells is 4-5 amino acids/s (Lodish & Jacobsen, 1972; Palmiter, 1974). The rate was corrected for our plants' lower growth temperature using the coefficient $Q_{10}$, which quantifies the rate of change of biochemical reactions when the temperature is varied by a factor of 10ºC,

$$Q_{10} = \left(\frac{R_2}{R_1}\right)^{\frac{10^\circ C}{T_2 - T_1}} \quad (2)$$

Where $R_2$ and $R_1$ are the rate of two reactions and $T_2$ and $T_1$ are the temperatures at which $R_2$ and $R_1$ are happening. Inverting this function and solving for the reaction $R_1$ occurring at the lower temperature $R_1$ results in,

$$R_1 = R_2 Q_{10}^{\frac{T_1 - T_2}{10^\circ C}} \quad (3)$$

Substituting the values R2=4.5 a.a./s, $T_2$=25.5ºC, $T_1$=21ºC, $Q_{10}$=2.5, yields an elongation rate estimate of 3 a.a./s at 21ºC (Piques et al., 2009). To select ribosomal loading, we assume that clock gene transcripts are in the large polysomal fraction and most timepoints are in the light, hence the value of 6 ribosomes per kb transcript is most relevant (Piques et al., 2009). The ORF length of each of the genes was obtained from TAIR10 (TAIR, 2016). These parameters were combined to estimate the rate of translation for each clock protein (Supplementary Table 1).

The simple model also requires a degradation rate estimate for each protein. Where decay curves had not been determined directly at the time (for evening complex proteins ELF3, ELF4 and LUX, as cited in the main text), minimum degradation rates were estimated as the rate required to match the falling phase of published protein timeseries (Nusinow et al., 2011). The timeseries from protein gel hybridisation images had not been quantified by the original authors. There are multiple limitations in extracting numerical timeseries from published images, *post hoc* using image analysis (ImageJ developers, n.d.). The 'Fermi problem' approach, however, is to make best use of any data that are available, as a precursor to acquiring more directly-relevant data. The resulting degradation rate estimates are listed in Supplementary Table 1.

#### 2. Deriving $K_d$ for promoter sequences by integrating Protein-Binding Microarray data and Surface Plasmon Resonance data.

In principle Protein-Binding Microarrays (PBMs) should capture the affinity landscape of a DNA-binding protein of interest for all DNA sequences. They should reflect the affinity obtained by other low-throughput, high-quality measurements of binding affinity, such as Surface Plasmon Resonance (SPR) data on $K_d$. We derived a linear, calibration model (Figure 5d) that links SPR data for the three binding sequences available for CCA1 (O'Neill et al., 2011) to PBM data on the same sequences for CCA1 (slope=-0.057, intercept=0.57, $r^2$=0.97, p-val=0.104) .

LUX affinity data were first reported for a single binding sequence (Silva et al., 2016). In this situation, the PBM data could still be extrapolated to estimate $K_d$ for LUX binding to other sequences but only by applying the same slope in the calibration model for LUX as for CCA1 (Figure 5e), a strong assumption but one that might be required in other cases. LUX $K_d$'s were reported for further binding sequences (Silva et al., 2020), allowing the same approach as for

CCA1 but returning a different slope in the linear regression (slope=-0.11, intercept=0.71, $r^2$=0.36, p-val=0.284) (Figure 5e). This model was used to estimate the $K_d$ for binding of the simulated EC to clock gene promoters (Table 2, Figure 4). The lower $r^2$ and different slopes for LUX and CCA1 likely derive from the technical differences between the $K_d$ assays, notably testing LUX binding by Electrophoresis Mobility Shift Assays (EMSA) rather than SPR.

#### 3. Using energy matrices obtained by Error-Model Averaging

In principle we could apply the PBM calibration described above to any sequence but it is not trivial to decide which collection of potential binding sequences is relevant. PBM experiments result in histograms of bound protein fluorescence where the E-score values from bound and unbound 8-mer sequences overlap to an unknown extent (Figure 5b) and technical variation also contributes. Selecting a single, statistical threshold would accept a subset of sequences as bound, above the threshold, and ignore sequences below the threshold. From the collection of bound sequences, Position Weight Matrices (PWM) could be derived (Godoy et al., 2011) to extrapolate the $K_d$ for any sequence, as we required. We avoided placing such a hard threshold on the overlapping distributions of bound and unbound signals, by inferring a biophysical model for binding, in the Error Model Average (EMA) approach (Kinney et al., 2007).

Briefly, this approach derives an energy matrix with an associated DNA-binding energy L as a classifier (Figure 5c). The original EMA was applied to raw protein-binding microarray data but only processed data in the form of enrichment scores (E-scores) could be obtained from the authors in our case. The continuous E-score distribution is discretised into $m$ bins. The sequences in each E-score bin can be split into bound ($x$=1) or unbound ($x$=0) according to a binding model. The model comprises a 4xL matrix $M$, where each position in the matrix represents the energy contribution of binding (in arbitrary units) by one of the four DNA bases A, T, G or C in a sequence. Each sequence position contributes additively to the total binding energy L. If the total energy is below a certain value, described below, then the sequence is classified as bound, otherwise as unbound. After sequence classification by the matrix, we compute the following likelihood, derived by Kinney

$$p(\{z_i\}|M) = \frac{\prod_{zx} \Gamma(c_{zx})}{\prod_{x} \Gamma(m - 1 + \sum_{z} c_{zx})}$$

$p(\{z_i\}|M)$ is the probability of a signal intensity given a matrix $M$, $c_{zx}$ is the number of sequences in the bin $z$ classified as bound $x$=1 or unbound $x$=0. Γ denotes the gamma function, the extended continuous case for the factorial, which simplifies the calculations during the inference process. A flat prior is assumed for the error model.

Kinney et al also described an MCMC Metropolis Hasting sampling scheme for inferring the maximum-likelihood parameters of the energy model, namely the columns of the matrix $M$ and the energy threshold for binding. Applying this approach to the E-score distribution yields an ensemble of energy matrices that captures the landscape of binding likelihood. The energy matrices effectively deconvolve the E-score distribution into distributions of bound and unbound 8-mer sequences.

#### 4. Assigning $K_d$ values to genes in the model

The promoter regions for clock genes that present a ChIP-seq signal (Ezer et al., 2017; Kamioka et al., 2016) indicating CCA1 (or LUX) binding were extracted from the TAIR10 genome annotation using Python scripts. The promoter region was considered from the transcription start site of the clock gene, spanning the intergenic region to the end of the next transcript annotated in the upstream gene. This region was scanned using the energy matrix derived using EMA, described above, as a classifier for bound or unbound sequences. Each 8-mer sequence classified as bound contributed its affinity $K_a$=1/$K_d$, from the calibration of E-scores into $K_d$

described in section 2 above. The full promoter's $K_a$ was calculated as the sum of the affinity contributions of each 8-mer, and the resulting total $K_a$ was inverted to give the total $K_d$ (Table 2).

As a simple validation for the approach, we tested whether these EMA-derived $K_d$'s for clock gene promoters were unusual in the genome. For each clock gene promoter, 1,000 regions of the same size as the promoter were sampled randomly from the genome. The total $K_d$ was calculated for each of the sample regions as above, ln-transformed and the z-score of each sample region was calculated as the S.D.-normalised difference from the mean $K_{d,}$, ($K_d$ - mean)/S.D (Table 2). The clock genes had unusually low EMA-derived $K_d$'s for CCA1 compared to other genomic sequences (z-scores 1.3 to 2.7), whereas their $K_d$'s for LUX were closer to the mean (z-scores 0.2-1.7). The specificity of EC binding *in vivo* might depend upon further sequences and their binding proteins, for example. Supplementary Table 5 notes possible future developments of this approach.

#### 5. Construction of transgenic plants with NanoLUC-tagged clock proteins

The genomic fragments of clock genes that were cloned to create the C-term fusions with NanoLUC-3FLAG-10His and the primers used are listed in Supplementary Tables 6 and 7. Equivalent fusions to the firefly LUC+ reporter were prepared in plasmid pGWB635.

##### 5.a Preliminary testing in protoplasts

We followed the suggestion of (Hansen & van Ooijen, 2016), to rapidly assess the functionality of the multiple NanoLUC reporter fusion constructs using a transient-expression system in leaf protoplasts of Col-0 plants (Supplementary Figure 3). Bioluminescent signals in this system indicate that the reporter gene is intact and also that appropriate RNA splicing has been maintained, in order to express these C-terminal reporter fusions. Protoplasts from plants of Col-0 grown under 16L:8D cycles for 4 weeks were transfected with 10µg of plasmid DNA

(Midi prep kit, Qiagen Ltd., Manchester) and tested under constant light, in a final dilution of 1:50 furimazine: Imagine Solution (Promega, Southampton, UK). LUC+ fusions were tested in Imagine Solution (1.2 mM Luciferin, W5 buffer, 1:1000 Amp and 5% FBS). Bioluminescence was measured in a Tristar $L^2$ luminometer (Berthold Technologies, Harpenden UK). Although mean reporter expression fell over several days (Supplementary Figure 3), it was clearly rhythmic and the relative phases of the first major peaks were as expected from the native promoters (*LHY*, *PRR9*, *PRR7* and *TOC1*). This method was valuable in validating complicated, synthetic constructs, consistent with the original authors' conclusion.

##### 5.b Mutant complementation

Functionality of the clock proteins with the C-terminal protein fusion was tested by genetic complementation of clock mutants or double mutants, in transgenic plants (Supplementary Figure 4).

Briefly, seeds were surface-sterilised in 0.01% Triton X-100, 5% bleach and washed three times in distilled water. 20 seeds of each independent line were sown in polypropylene collars made from 1.5 ml microcentrifuge tubes, on solid media containing 1/2x Murashige and Skoog salts (Duchefa) pH 5.8, 1.2% agar. Seed were stratified for 3 days and then transferred to 120 µmol $m^{-2}s^{-1}$ cool white fluorescent light under 12L:12D cycles at 21ºC. After one week the plants were transferred to imaging conditions under monochromatic blue and red LED light with a total 50 µmol $m^{-2}s^{-1}$, 12L:12D photoperiod, at 21ºC. For firefly LUC assays, plants were sprayed with 5 mM luciferin, 0.01% Triton X-100. After one day in these conditions, image acquisition every 1.5h started using Hamamatsu OrcaII digital cameras operating at -75ºC (model C4742-98, Hamamatsu Photonics, Hamamatsu City, Japan), with constant lighting between images under the control of the manufacturer's Wasabi software. The raw images were imported into ImageJ64 software (ImageJ developers, n.d.) and bioluminescent signals were analysed in circular regions of interest.

The short-period phenotype of the *lhy-1/cca1-11* CCA1p:LUC line could be complemented to wild-type period either by LHY or by CCA1 fusion protein (Supplementary Table 8), with each at similar levels (Table 1, Figure 6a, Supplementary Figure 5). Some transformants for each transgene showed even longer periods than the wild type, suggestive of transgene over-expression (Supplementary Figure 4a). Whereas the double-mutant parents showed the expected, hypocotyl shortening compared to wild type seedlings, the hypocotyls of the CCA1-complemented seedlings were appreciably longer than the hypocotyls of LHY-complemented seedlings, similar to observations for CCA1-overexpressing plants (Lu et al., 2012). The short period of the *toc1* mutant parent and the long period of the *prr7prr9* double mutant were also restored to wild-type period (Supplementary Table 8 shows the lines selected) or even longer by the TOC1 reporter fusion protein, or shorter by the PRR7 fusion, suggestive of overexpression (Supplementary figures 4b, 4c). Quantitative complementation of the *prr7* single mutant was infeasible, because the single mutant has such a weak period phenotype. Transgenic *elf3-1* mutants containing the ELF3p:ELF3-NL3F10H construct were selected for complementation of the mutant's long-hypocotyl phenotype, under 8L:16D cycles of cool white fluorescent light at 22ºC, retaining three transgenic lines that showed hypocotyl lengths close to wild-type seedlings (Supplementary Figure 4g). Transgenic lines carrying the LUX fusion complemented the *lux* mutant's period defect close to the wild-type period, with few partially-complemented or over-complemented lines. Rather, most lines appeared close to the wild-type period (Supplementary Figure 4d). We speculate that the dose-dependence of period on LUX expression level might differ from that of the other clock proteins tested. A separate test of hypocotyl elongation revealed period-complemented lines where the *lux* mutant's long hypocotyl was not restored to wild type (not shown), including the line 21.2 that was tested in extracts (Supplementary Figures 4e, 5c). Line 9.2 used for *in vivo* assays (Supplementary Figure 4f, 4g) complemented both period and hypocotyl phenotypes. This complicates the inference of native LUX protein expression levels. The low levels quantified in extracts of line 21.2 restored only period but not hypocotyl regulation in the transgenic mutant host, whereas the higher levels estimated *in vivo* from line 9.2 complemented both phenotypes (Supplementary Figure 4).

### Supplementary Information References

Ezer, D., Jung, J. H., Lan, H., Biswas, S., Gregoire, L., Box, M. S., Charoensawan, V., Cortijo, S., Lai, X., Stockle, D., Zubieta, C., Jaeger, K. E., & Wigge, P. A. (2017). The evening complex coordinates environmental and endogenous signals in Arabidopsis. *Nature Plants, 3*, 17087. https://doi.org/10.1038/nplants.2017.87

Flis, A., Fernandez, A. P., Zielinski, T., Mengin, V., Sulpice, R., Stratford, K., Hume, A., Pokhilko, A., Southern, M. M., Seaton, D. D., McWatters, H. G., Stitt, M., Halliday, K. J., & Millar, A. J. (2015). Defining the robust behaviour of the plant clock gene circuit with absolute RNA timeseries and open infrastructure. *Open Biology, 5*(10). https://doi.org/10.1098/rsob.150042

Godoy, M., Franco-Zorrilla, J. M., Pérez-Pérez, J., Oliveros, J. C., Lorenzo, Ó., & Solano, R. (2011). Improved protein-binding microarrays for the identification of DNA-binding specificities of transcription factors. *The Plant Journal*, *66*(4), 700–711. https://doi.org/10.1111/j.1365-313X.2011.04519.x

Hansen, L. L., & van Ooijen, G. (2016). Rapid Analysis of Circadian Phenotypes in Arabidopsis Protoplasts Transfected with a Luminescent Clock Reporter. *JoVE (Journal of Visualized Experiments), 115*, e54586. https://doi.org/10.3791/54586

ImageJ developers. (n.d.). *ImageJ.net*. Retrieved August 23, 2024, from https://imagej.net/ij/download.html

Kamioka, M., Takao, S., Suzuki, T., Taki, K., Higashiyama, T., Kinoshita, T., & Nakamichi, N. (2016). Direct Repression of Evening Genes by CIRCADIAN CLOCK-ASSOCIATED1 in the Arabidopsis Circadian Clock. *The Plant Cell*, *28*(3), 696–711. https://doi.org/10.1105/tpc.15.00737

Kinney, J. B., Tkacik, G., & Callan, C. G. (2007). Precise physical models of protein-DNA interaction from high-throughput data. *Proceedings of the National Academy of Sciences of the USA*, *104*(2), 501–506. https://doi.org/10.1073/pnas.0609908104

Lodish, H. F., & Jacobsen, M. (1972). Regulation of Hemoglobin Synthesis. *Journal of Biological Chemistry*, *247*(11), 3622–3629. https://doi.org/10.1016/S0021-9258(19)45186-7

Lu, S. X., Webb, C. J., Knowles, S. M., Kim, S. H., Wang, Z., & Tobin, E. M. (2012). CCA1 and ELF3 Interact in the control of hypocotyl length and flowering time in Arabidopsis. *Plant Physiology*, *158*(2), 1079–1088. https://doi.org/10.1104/pp.111.189670

Nusinow, D. A., Helfer, A., Hamilton, E. E., King, J. J., Imaizumi, T., Schultz, T. F., Farré, E. M., & Kay, S. A. (2011). The ELF4–ELF3–LUX complex links the circadian clock to diurnal control of hypocotyl growth. *Nature*, *475*(7356), 398–402. https://doi.org/10.1038/nature10182

O'Neill, J. S., van Ooijen, G., Le Bihan, T., & Millar, A. J. (2011). Circadian clock parameter measurement: Characterization of clock transcription factors using surface plasmon resonance. *Journal of Biological Rhythms*, *26*(2), 91–98. https://doi.org/10.1177/0748730410397465

Palmiter, R. D. (1974). Differential Rates of Initiation on Conalbumin and Ovalbumin Messenger Ribonucleic Acid in Reticulocyte Lysates. *Journal of Biological Chemistry*, *249*(21), 6779–6787. https://doi.org/10.1016/S0021-9258(19)42126-1

Piques, M., Schulze, W. X., Hohne, M., Usadel, B., Gibon, Y., Rohwer, J., & Stitt, M. (2009). Ribosome and transcript copy numbers, polysome occupancy and enzyme dynamics in Arabidopsis. *Molecular Systems Biology*, *5*, 314. https://doi.org/10.1038/msb.2009.68 msb200968 [pii]

Silva, C. S., Lai, X., Nanao, M., & Zubieta, C. (2016). The Myb domain of LUX ARRHYTHMO in complex with DNA: Expression, purification and crystallization. *Acta Crystallographica Section F: Structural Biology Communications*, *72*(5), 356–361. https://doi.org/10.1107/S2053230X16004684

Silva, C. S., Nayak, A., Lai, X., Hutin, S., Hugouvieux, V., Jung, J.-H., López-Vidriero, I., Franco-Zorrilla, J. M., Panigrahi, K. C. S., Nanao, M. H., Wigge, P. A., & Zubieta, C. (2020). Molecular mechanisms of Evening Complex activity in Arabidopsis. *Proceedings of the National Academy of Sciences*, *117*(12), 6901–6909. https://doi.org/10.1073/pnas.1920972117

TAIR. (2016). *Gene Annotation Data at TAIR*. https://v2.arabidopsis.org/portals/genAnnotation/gene_structural_annotation/annotation_data.jsp

Urquiza-García, U., & Millar, A. J. (2021). Testing the inferred transcription rates of a dynamic, gene network model in absolute units. *In Silico Plants*, *3*(2), diab022. https://doi.org/10.1093/insilicoplants/diab022
